# Establishment of a novel non-integrated human pluripotent stem cell-based gastruloid model

**DOI:** 10.1101/2023.06.28.546720

**Authors:** Gege Yuan, Jiachen Wang, Zhaode Liu, Mengqi Chen, Pinmou Zhu, Hao Zhang, Zhibin Hu, Yiqiang Cui, Yan Yuan, Jiahao Sha

## Abstract

Embryo loss and pregnancy disorders are prevalent worldwide, with both conditions critically associated with dysfunctioning gastrulation processes. Gastrulation and post-gastrulation organogenesis are crucial stages of embryonic development that establish the blueprint for body part formation. These processes involve the sequential generation of three germ layer cells and primordial germ cells, as well as the assembly of the precursor tissues for body parts. However, due to ethical limitations associated with studying human embryogenesis, a more detailed understanding of gastrulation and post-gastrulation organogenesis remains elusive. To ensure that the knowledge obtained from gastruloids is biologically meaningful and clinically relevant, it is critical to create high-fidelity human embryo models that closely mimic embryogenesis *in vivo*. Here, we developed a two-stage derivation gastruloids *in vitro* based on human pluripotent stem cells. Morphological tracking mimicks the developmental processes of models from Carnegie Stage 4 (CS4) to early CS7. Our gastruloids exhibit key structures characteristic of human embryos, including amniotic cavity, embryonic disc, primitive streak, primary yolk sac, secondary yolk sac, and blood islets. Comparison of our cell lineage development maps showed that gastruloids closely resembled human natural CS7 gastrula. Our gastruloids exhibited transcriptional characteristics that mimicked the molecular pathways observed in natural embryos development. Importantly, we found that in our model, extraembryonic mesoderm originates from the yolk sac and primordial germ cells originate from the posterior epiblast of the embryonic disc. Moreover, we found that thalidomide affects the differentiation of three germ layer cells, resulting in the arrest of human gastruloid development. In conclusion, by establishing a human gastruloid, we were able to gain valuable insights into the mechanisms responsible for human gastrulation and shed light on the causes of early embryo loss and pregnancy disorders.

## Introduction

Birth defects affect eight million newborns every year globally. Approximately 8% of global mortality below 5-year age is attributed to birth defects^1^. Numerous agents can cause malformation in the developing embryo, including drugs, chemicals, and viruses. In addition, faulty genes and chromosomal abnormalities can also be teratogenic. Large population-based multicenter case-control studies of birth defects have been used to identify teratogens responsible for he field of birth defect epidemiology^2,3^. However, teratogens identified to date only account for the tip of the iceberg, and efficient screening methods need to be established to identify more teratogens^4^.

Human embryos start to implant in the uterus from Carnegie Stage 4 (CS4), and peri-implantation embryos differentiate to form the amniotic cavity, yolk sac, and bilaminar embryonic disc. This process is followed by gastrulation, with post-gastrulation organogenesis including critical phases of embryonic development, during which the blueprint of development is established through the ordered generation of three germ layer cells and primordial germ cells (PGCs), as well as the assembly of the precursor tissues for body parts^5^. However, little is known about the biology of human gastrulation, as clinical samples are difficult to obtain, and ethical concerns limit the study of human embryogenesis. An immediate challenge is to create human gastruloid models of high fidelity truthfully representing embryogenesis *in vivo* to ensure that the knowledge gained from these studies is biologically meaningful and clinically relevant. Human stem cell-based models have been created to recapitulate some aspects of early mammalian development in a modular manner, such as micropattern, amniotic organoids, asymmetric division organoids, and gastruloids^6–9^.While these models have considerably contributed to a more detailed understanding of human gastrulation, most of the existing models are assembled using by different cell types, or combined with new materials and bioengineering, and have not yet been able to create models faithfully mimicking natural human gastrula.

In this study, we constructed an atlas for the development of human gastrula through establishment of a non-integrated gastruloid utilizing human pluripotent stem cells. This model provides a high-throughput screening platform for the discovery of teratogens that cause disorders associated with embryonic development.

## Results

### Establishment and characterization of human gastruloids

In order to establish human gastruloids *in vitro*, we employed a two-stage derivation process utilizing two pluripotent stem cell (hPSC) lines, namely DYR0100 hiPSC (human induced pluripotent stem cell, hereafter referred to as hiPSC), and H1 hESC (human embryonic stem cell, hereafter referred to as hESC). hPSC contains pluripotent founder cell (hPFC) that shares transcriptional features with the hypoblast and maintains the pluripotency^10^. We mixed G-1^TM^ plus, E8 medium (maintained the pluripotency) and RACL (induced the hypoblast^11^), at a 2:1:1 ratio to form the first-stage medium. This medium was then employed to induce the hypoblast while preserving partial pluripotency, forming aggregates of hPFC. Basic fibroblast growth factor (bFGF) and bone morphogenetic protein (BMP4) have previously been shown to induce differentiation of the amnion and gastrulation^12^. In the second-stage, we combined G-2^TM^ plus, E8 medium and EbB (E6 medium, bFGF, and BMP4) at a ratio of 2:1:1. The aggregates gradually completed amnion formation, primordial germ cell (PGC) specification and gastrulation, finally formed gastruloids (Fig. 1a).

**Fig. 1.**
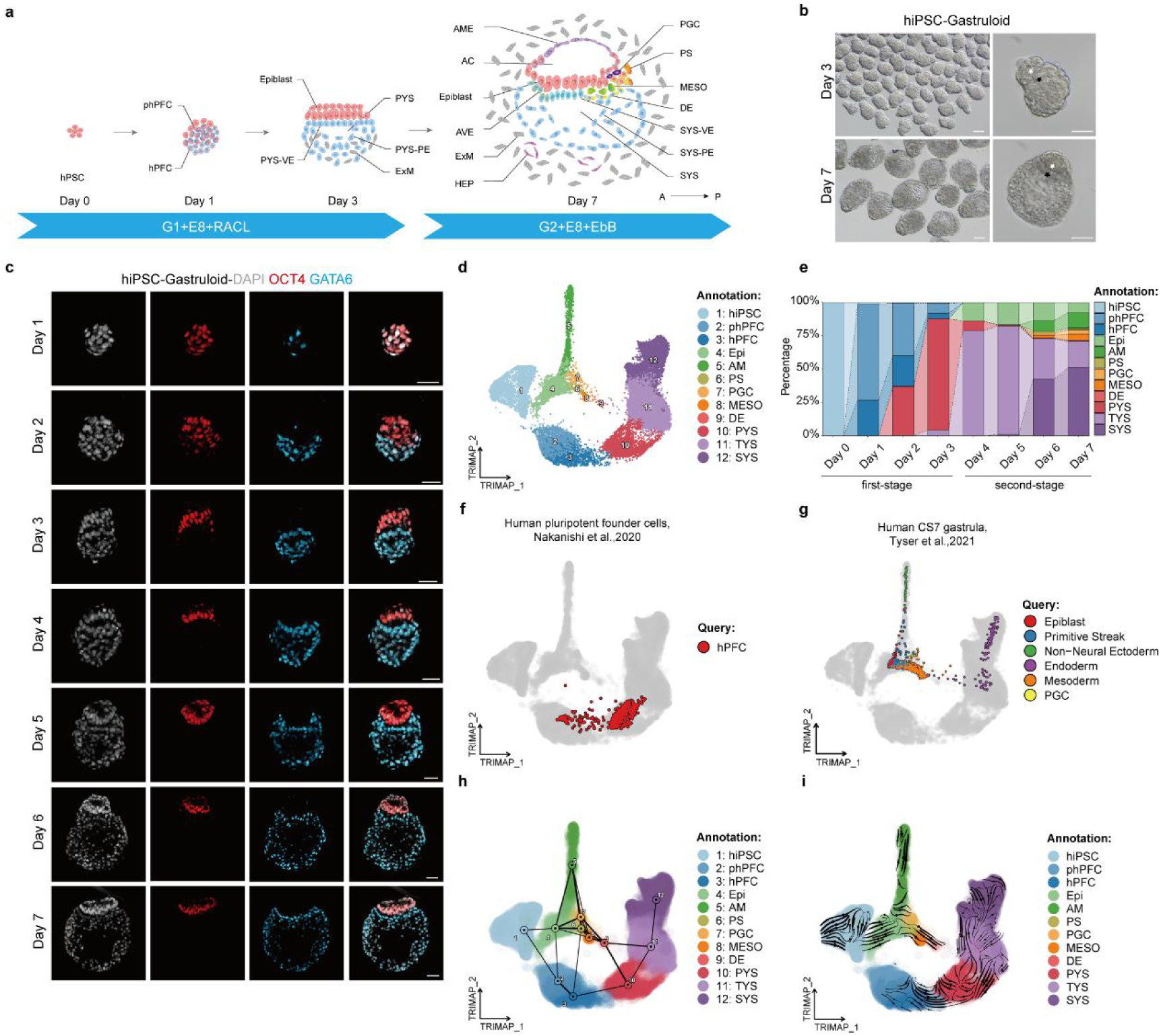
Establishment and characterization of human gastruloids. (a) Schematic of human gastruloids two-stage induction. hPSC, human pluripotent stem cell; hPFC, human pluripotent founder cell; phPFC, pre-hPFC; PYS, primary yolk sac; PYS-VE, PYS-visceral endoderm; PYS-PE, PYS-parietal endoderm; ExM, extraembryonic mesoderm; AME, amniotic epithelium; AC, amniotic cavity; AVE, anterior visceral endoderm; PGC, primordial germ cell; PS, primitive streak; MESO, mesoderm; DE, definitive endoderm; SYS, secondary yolk sac; SYS-VE, SYS-visceral endoderm; SYS-PE, SYS-parietal endoderm; HEP, hematopoietic endothelial progenitor cell; A, anterior axis; P, posterior axis. (b) Brightfield images of representative the human gastruloid (based on hiPSC) on day 3 and 7. White asterisks, epiblast. Black asterisks, hypoblast. Scale bars, 50 μm (left) and 100 μm (right). (c) Immunofluorescence (IF) staining for epiblast marker OCT4 (red) and hypoblast marker GATA6 (blue) of the human gastruloid (based on hiPSC) from day 1 to 7. Nuclei were counterstained with DAPI (grey). Scale bar, 50 μm. (d) TriMap showing the 12 major cell types. TYS, transitional yolk sac. (e) Bar plot showing the proportions of 12 clusters in each sampling day. (f) TriMap projection of hPFC from Nakanishi et al.^10^ onto this study (grey). (g) TriMap projection of human CS7 gastrula dataset^13^ onto day 7 of this study (grey). (h) PAGA analysis overlaid on TriMap. he threshold for connection of cell types was set to 0.035. (i) RNA velocity plotted on TriMap.

We aliquoted 50-75 hiPSCs per microwell and cultured these cells for 3 days in the first-stage medium. We found that these cells were highly proliferative and gradually formed spherical asymmetric aggregates (Fig. 1b, Extended Data Fig. 1a). Using immunofluorescence (IF) staining, we observed the emergence of OCT4^+^ GATA6^+^ cells within the aggregates on day 1, which suggested that hiPSCs were differentiating into hPFCs (Fig. 1c). On day 2, GATA6^+^ hPFCs proliferated and migrated to one side, leading to the separation of OCT4^+^ cells and GATA6^+^ cells in aggregates (Fig. 1c). This separation resulted in the establishment of polarity and the specializition into the hypoblast. On day 3, the hypoblast surrounded a cavity to form the rudimentary primary yolk sac (PYS) (Fig. 1c, Extended Data Fig. 1b). Next, we proceeded with the second-stage induction and the addition of bFGF and BMP4 facilitated the formation of gastruloids (Fig. 1b, Extended Data Fig. 1a). From day 4 to 5, the OCT4^+^ epiblast underwent epithelialization and cavitation, culminating in the formation of the amniotic cavity (Fig. 1c). As the amniotic cavity expanded, the embryonic disc formed, which was composed of a tall columnar epiblast (OCT4^+^) at the bottom of the amniotic cavity and a short columnar hypoblast (GATA6^+^) at the top of the PYS. From day 6 to 7, the hypoblast separated the PYS into several small sacs which gradually fused to form a secondary yolk sac (SYS) (Fig.1c). In particular, the embryonic disc cells gradually proliferated and migrated, suggested that gastrulation emerged in these aggregates (Fig. 1c, Extended Data Fig. 1c, Supplementary Movie 1). On the basis of these observations, we concluded that we successfully obtained the gastruloids. During the entire induction process, the gastruloids stably proliferated (Extended Data Fig. 1d). Using these morphological criteria, we then quantified the efficiency of gastruloid formation as approximately 48% (Extended Data Fig. 2e). hESC formed such structures with lower efficiency (around 30%) (Extended Data Fig. 2a-e). These results clearly demonstrated that we established a non-integrated hPSC-based gastruloid through a three-dimensional (3D) two-stage derivation protocol.

**Fig. 2.**
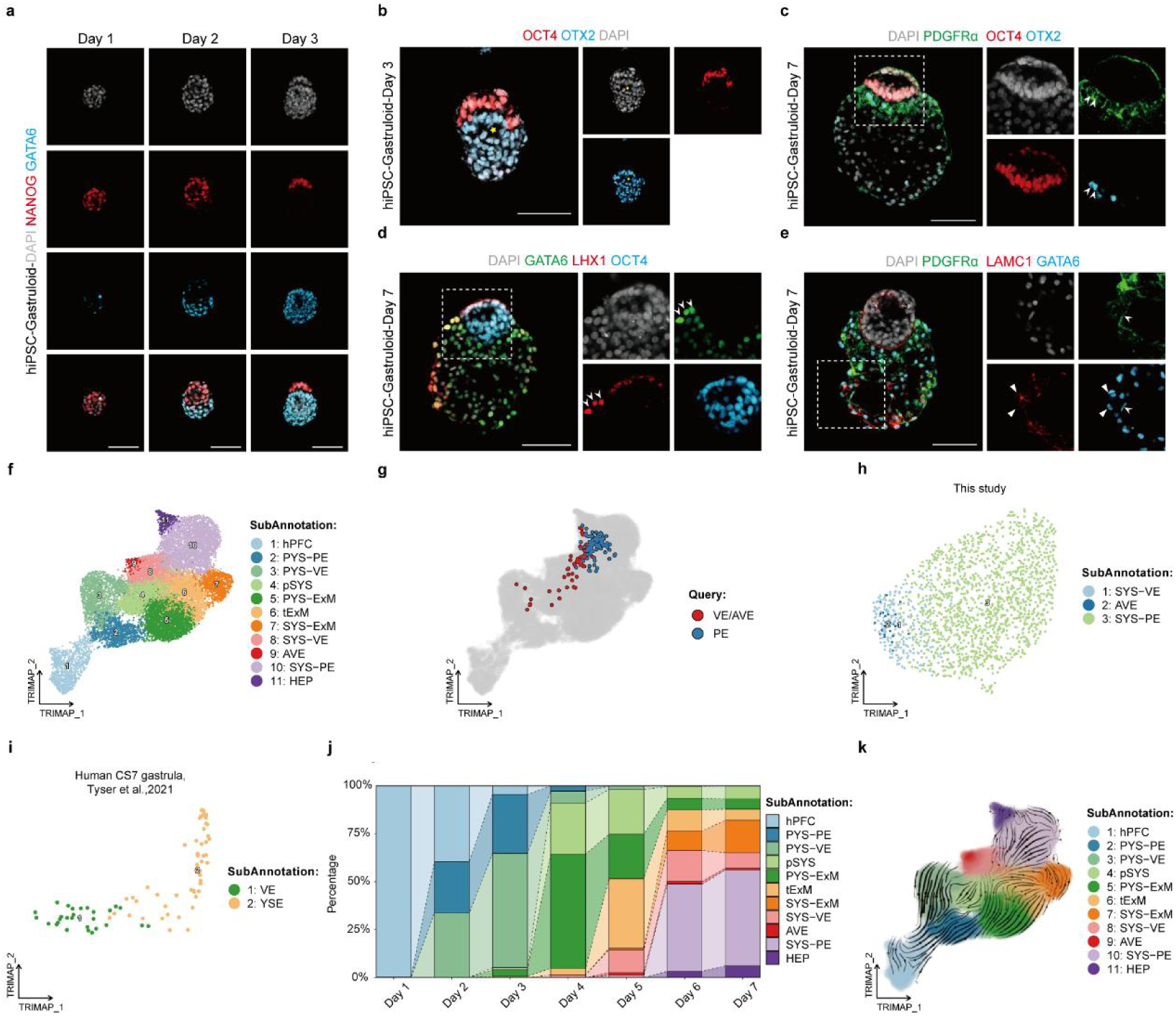
The development and characterization of the yolk sac in gastruloids. (a) IF staining for epiblast marker NANOG (red) and hypoblast marker GATA6 (blue) of the human gastruloid (based on hiPSC) from day 1 to 3. Nuclei were counterstained with DAPI (grey). Scale bars,100 μm. (b) IF staining for epiblast marker OCT4 (red) and hypoblast/ anterior visceral endoderm marker OTX2 (blue) of the human gastruloid (based on hiPSC) on day 3. Nuclei were counterstained with DAPI (grey). Yellow asterisk, primary yolk sac. Scale bar, 100 μm (c) IF staining for epiblast marker OCT4 (red), hypoblast/anterior visceral endoderm marker OTX2 (blue) and secondary yolk sac marker PDGFRα (green) of the human gastruloid (based on hiPSC) on day 7. Nuclei were counterstained with DAPI (grey). White arrowhead, anterior visceral endoderm. Scale bar, 100 μm. (d) IF staining for epiblast marker OCT4 (blue), anterior visceral endoderm marker LHX1 (red) and hypoblast marker GATA6 (green) of the human gastruloid (based on hiPSC) on day 7. Nuclei were counterstained with DAPI (grey). White arrowhead, anterior visceral endoderm. Scale bar, 100 μm. (e) IF staining for hypoblast marker GATA6 (blue), secondary yolk sac marker PDGFRa (green) and extraembryonic mesoderm marker LAMC1 (red) of the human gastruloid (based on hiPSC) on day 7. Nuclei were counterstained with DAPI (grey). White arrowhead, secondary yolk sac. White triangle arrow, extraembryonic mesoderm. Scale bar, 100 μm. (f) TriMap showing 11 subtypes of hypoblast derivatives. hiPSC, human induced pluripotent stem cell; hPFC, human pluripotent founder cell; PYS-PE, primary yolk sac -parietal endoderm; PYS-VE, PYS-visceral endoderm; pSYS, precursor of secondary yolk sac; SYS-VE, SYS-visceral endoderm; SYS-PE, SYS-parietal endoderm; AVE, anterior visceral endoderm; HEP, haemato-endothelial progenitor; PYS-ExM, PYS-extraembryonic mesoderm, tExM, transition state of ExM. (g) TriMap projection of the VE/AVE and PE from the integrated human embryonic reference datasets onto this study (grey). (h-i) TriMaps of SYS-VE, SYS-PE and AVE in gastruloids on day 7 (h) integrated with human CS7 gastrula (i). (j) Bar plot showing the proportions of 11 subtypes in each sampling day. (k) The RNA velocity analysis of the hypoblast differentiation.

To investigate the composition and cell lineage differentiation of gastruloids, single-cell RNA-sequencing (scRNA-seq) was performed on gastruloids from hiPSCs at eight time points (every 24 hours) using the 10x Genomics platform, and 27,737 cells passed the quality control. We integerated these datasets using Scanorama and reduced the dimensions using TriMap (triplet manifold approximation and projection) to generate the development map of gastruloids (Fig. 1d). We found that our gastruloids formed 12 major cell types, as based on expression patterns of key lineage markers, namely hiPSC, pre-hPFC (phPFC), hPFC, epiblast (Epi), amnion (AM), PS, PGC, mesoderm (MESO), definitive endoderm (DE), PYS, transitional yolk sac (TYS), and SYS (Fig. 1d, Extended Data Fig. 3a-c, Supplementary Table 1). Additionally, we found that the gastruloids developed in an orderly manner. During the first-stage, development of the hypoblast occurred and during the second-stage, embryonic disc, amnion, PGC, gastrulation, SYS and their derivatives appeared (Fig. 1e, Extended Data Fig. 3a). When we performed cell cycle analysis, we found that both the PYS and SYS cells were predominantly arrested in G1 phase, while the Epi and PS cells exhibited more active proliferation, as indicated by the majority of cells in G2M phase (Extended Data Fig. 4a).

We next focused on the process of hiPSC transforming into hPFC in gastruloids. When we projected the reported single-cell dataset of hPFC^10^ onto our cell populations in the first-stage, we found that their hPFC exhibited similar traits to our hPFC and PYS (Fig. 1f). Moreover, this transformation process was accompanied by a decrease in the expression of pluripotency-related markers and an increase in the expression of hPFC-specific markers (Extended Data Fig. 4b). Next, we compared with the dataset of human CS7 gastrula^13^. We found that our gastruloids on day 7 were transcriptionally similar to human CS7 gastrula (Fig. 1g). The developmental trajectories of the cell lineages in gastruloids inferred by single-cell RNA velocity (scVelo) and partition-based graph abstraction (PAGA) were consistent with the developmental relationships between cell types (Fig. 1h, i).

Together, the results suggested that our gastruloid simulated cell lineage development from hPFC to gastrula, both in terms of morphology and transcription.

### Yolk sac development

To reveal the origin and development of the human yolk sac, we first traced the specialization of hypoblast in gastruloids by IF. From day 1 to 3, we observed the emergence of GATA6 expression within a subpopulation of NANOG^+^ cells, upon which the GATA6^+^ cells separated from the NANOG^+^ cells to form the “salt and pepper” pattern^14^, with the hypoblast proliferating and migrating downward (Fig. 2a). This observation suggested that the development of the hypoblast had already occurred. Moreover, at this stage, we found that OTX2 was diffusely expressed throughout the hypoblast (Fig. 2b), indicating that the embryonic axis had not been established ^15,16^. In the second-stage, the PYS expanded further. On day 6, GATA6^+^ cells located below OCT4^+^ epiblast arranged to form visceral endoderm (VE) and the remaining GATA6^+^ cells formed the SYS (Fig. 1c). On day 7, a group of OTX2^+^ or LHX1^+^ cells asymmetrically positioned themselves at the anterior of the embryonic disc (Fig. 2c, d, Supplementary Movie 2), indicating that anterior visceral endoderm (AVE) appeared and anterior-posterior (AP) axis established^15^. Moreover, the SYS composed of PDGFRα^+^ cells continued to enlarge (Fig. 2c, e), with being surrounded by LAMC1^+^ extraembryonic mesoderm (ExM)^17^ (Fig. 2e). To verify the reproducibility of the hypoblast development in our gastruloids, we performed IF staining on hESC-based gastruloids on day 7 (Extended Data Fig. 5a, b). We found that hESC-based gastruloids exhibited AVE, SYS, and ExM, thereby faithfully recapitulating the hypoblast development in gastruloids.

To investigate the molecular properties of cell lineage development from PYS to SYS in gastruloids, we re-clustered the hypoblast and its derivatives, defining 11 cell types based on the expression of known markers and proliferation characteristics^18^ (Fig. 2f, Extended Data Fig. 6a-c). We categorized these cell types into hPFC, parietal endoderm of PYS (PYS-PE), visceral endoderm of PYS (PYS-VE), Pre-SYS (pSYS), SYS-VE, SYS-PE, AVE, and haemato-endothelial progenitors (HEP), and progressively maturing ExM including PYS-ExM, Tran-ExM (tExM), and SYS-ExM (Fig. 2f). Next, we projected the published data of *in vitro* cultured (IVC) human embryos^19^, human pre-implantation embryos^20^, and IVC human post-implantation embryos^15^ onto our data, verified that the overall transcriptome differences between PE and VE of gastruloids were consistent with the reported data (Fig. 2g, Extended Data Fig. 6d-g). To compare the similarity of the hypoblast and its derivatives between gastruloids and natural gastrula, we integrated the corresponding cell types in our data on day 7 with that of human CS7 gastrula^13^. We found that the SYS-VE and SYS-PE in the gastruloids were significantly similar to VE and the yolk sac endoderm (YSE) of human CS7 gastrula, respectively (Fig. 2h, i, Extended Data Fig. 6h, i). Further comparison with single-cell transcriptomes derived from marmoset embryos^21^ showed that VE/AVE and SYS-PE in the gastruloids exhibited close correlation in gene expression to respectively the VE and SYS of marmosets (Extended Data Fig. 6j, k, Supplementary Table 3). However, the ExM in gastruloids displayed characteristics of both the endoderm and mesoderm of marmosets (Extended Data Fig. 6j, k, Supplementary Table 3). In addition, we found that the PYS were gradually replaced by the subsequent SYS, HEP and ExM, which increased in proportion (Fig. 2j).

The mammalian yolk sac originates from the hypoblast, an extraembryonic lineage stemming from the early inner cell mass of the blastocyst^22^. Our analysis of the dynamic features along lineages indicated that a key transitional point from epiblast to hypoblast was present in the downregulation of epiblast markers and upregulation of endoderm markers (Extended Data Fig. 7a, b). We also observed that GATA6, SOX17, and GATA4 were sequentially upregulated in gastruloids (Extended Data Fig. 7b), consistent with the transcriptional program governing primate hypoblast specification^17,23–25^. Using IF staining, we observed more GATA6^+^ cells than GATA4^+^ cells in our gastruloids on day 2 (Extended Data Fig. 7c). This suggested that GATA6 was expressed prior to GATA4 during hypoblast specification, which was consistent with the transcriptome results. Genes related to positive regulation of TGF-β and Wnt pathways were found upregulated in pSYS, suggesting selective activation of these pathways during hypoblast differentiation (Extended Data Fig. 7d).

When we conducted both scVelo and PAGA analysis, we found that PYS-VE transformed into pSYS and further developed into SYS-VE, SYS-PE, and HEP, while PYS-PE changed into PYS-ExM and directly progressed through tExM to develop into SYS-ExM. Additionally, pSYS also differentiated into tExM, before ultimately developing into SYS-ExM (Fig. 2k, Extended Data Fig. 8a, b). To identify branching points between trajectories, we separately calculated the differential developmental genes (DDGs) for each pseudotime trajectory (Extended Data Fig. 8c, d). We found that C4 DDGs were enriched with genes related to the mesoderm. In comparison, C5 DDGs such as *APOA2, APOC3* and other carrier protein family members correspond to the well-established functions of the human YSE. These functions included the absorption of nutrients from the exocoelomic cavity and the transportation of these nutrients to the developing embryo^18,26^ (Extended Data Fig. 8d, Supplementary Table 2). We noted that the subsequent developmental trajectory of PYS-PE displayed a stronger overall proliferation level than PYS-VE^18^ (Extended Data Fig. 8e).

Taken together, these findings clearly demonstrated that the morphology and molecular properties underlying the formation of SYS as well as the origin and development of ExM.

### Fate determination of epiblast development

In order to gain insight into the fate determination of the epiblast, we tracked the developmental process of the epiblast during the second-stage using IF staining. We found that on day 4, the OCT4^+^ epiblast cells underwent cavitation, resulting in the rise of the pre-amniotic cavity (Fig. 1c). This rudimentary structure had developed into the amniotic cavity on day 5 (Extended Data Fig. 9a). On day 6, SOX2^-^ ISL1^+^ amniotic precursor cells emerged at the apical region of the EZRIN^+^ amniotic cavity (Fig. 3a). The appearance of SOX2^-^ OCT4^+^ TBXT^+^ gastrulating cells within the embryonic disc, served as a clear indicator that gastruloids had formed (Fig. 3b). When we further tracked amniotic development and gastrulation, we found that on day 7, the amniotic epithelial cells transformed into SOX2^-^ ISL1^+^ / SOX2^-^ AP2α^+^ KRT7^+^ squamous epithelial cells (Fig. 3c, Extended Data Fig. 9b). OCT4^+^ TBXT^+^ cells migrated and formed either TBXT^+^ EOMES^+^ or TBXT^+^ MIXL1^+^ mesendoderm, which finally developed into the TBXT^-^ EOMES^+^ DE or TBXT^-^ MIXL1^+^ nascent mesoderm (NaM) (Fig. 3d, Extended Data Fig. 9c, Supplementary Movie 3). We also found that VIM^+^ advanced mesoderm (AdM) derived from PS on day 7 were present in the gastruloid (Extended Data Fig. 9d). On day 6, we identified PGCs, which were subsequently found to be AP2γ^+^ NANOG*^+^*SOX17^+^ / AP2γ^+^ BLIMP1^+^SOX17^+^ PGCs in both the amnion and epiblast on day 7 (Fig. 3e, f, Extended Data Fig. 9e). To verify the reproducibility of the epiblast development in our gastruloids, we performed IF staining on hESC-based gastruloids on day 7 (Extended Data Fig. 9f-h). We found that hESC-based gastruloids exhibited amnion development, gastrulation, and PGCs specification, thereby faithfully recapitulating the epiblast development in gastruloids. To elucidate the molecular properties underlying the developmental process of the epiblast in gastruloids, we re-clustered the epiblast and its derivatives and divided them into 18 subtypes based on distinct molecular characteristics, namely hiPSC, phPFC, hPFC, PYS, Epi, pre-PGC (pPGC), primitive streak anlage-epiblast (PSA-Epi), PS, PGC, DE-1, DE-2, NaM, emergent mesoderm (EmM), AdM, HEP, amnion anlage-epiblast (AMA-Epi), AM-Early (AM-E), and AM-Late (AM-L) (Fig. 3g, Extended Data Fig. 10a-c). To characterize the features of Epi, AMA-Epi, PSA-Epi and pPGC, we performed a similarity analysis among these cell types and analyzed the differences in their trait transcriptomes using differentially expressed genes (DEGs) (Extended Data Fig. 10d, e, Supplementary Table 1). We observed that Epi, AMA-Epi, PSA-Epi, and pPGC exhibited notable similarities in specific aspects of their transcriptomes, while they simultaneously preserved their unique transcriptional features. As indicated by scVelo analysis, we found that the hPFC, derived from hiPSC, generated the PYS during the first-stage, and the remaining phPFC gave rise to the epiblast. The epiblast gradually diverged into AMA-Epi, PSA-Epi, and pPGC, ultimately contributing to the formation of amnion, mesoderm, DE, and PGCs (Fig. 3h, i). We next integrated the composition of the epiblast derivatives in the gastruloids on day 7 with that of human CS7 gastrula^13^ (Extended Data Fig. 11a, b). We found that our gastruloids contained most of the cell types present in human CS7 gastrula (Fig. 3j, k). To further assess the reliability of our data, we performed a similarity analysis with transcriptome data from the marmoset epiblast and its derivatives. We found that the epiblast in our gastruloids displayed the highest similarity with the epiblast of marmoset. Furthermore, Epi-derived PGCs and amnion exhibited high similarity with the corresponding cell types of marmoset (Extended Data Fig. 11c, d, Supplementary Table 3). Next, we projected the data from *in vitro* amnion-like cell populations^27^ onto our data and performed the similarity comparison. We found that the development of amnion in the gastruloid was similar to two developmental waves of amnion occurrence in human embryos (Extended Data Fig. 11e-g, Supplementary Table 3).

**Fig. 3.**
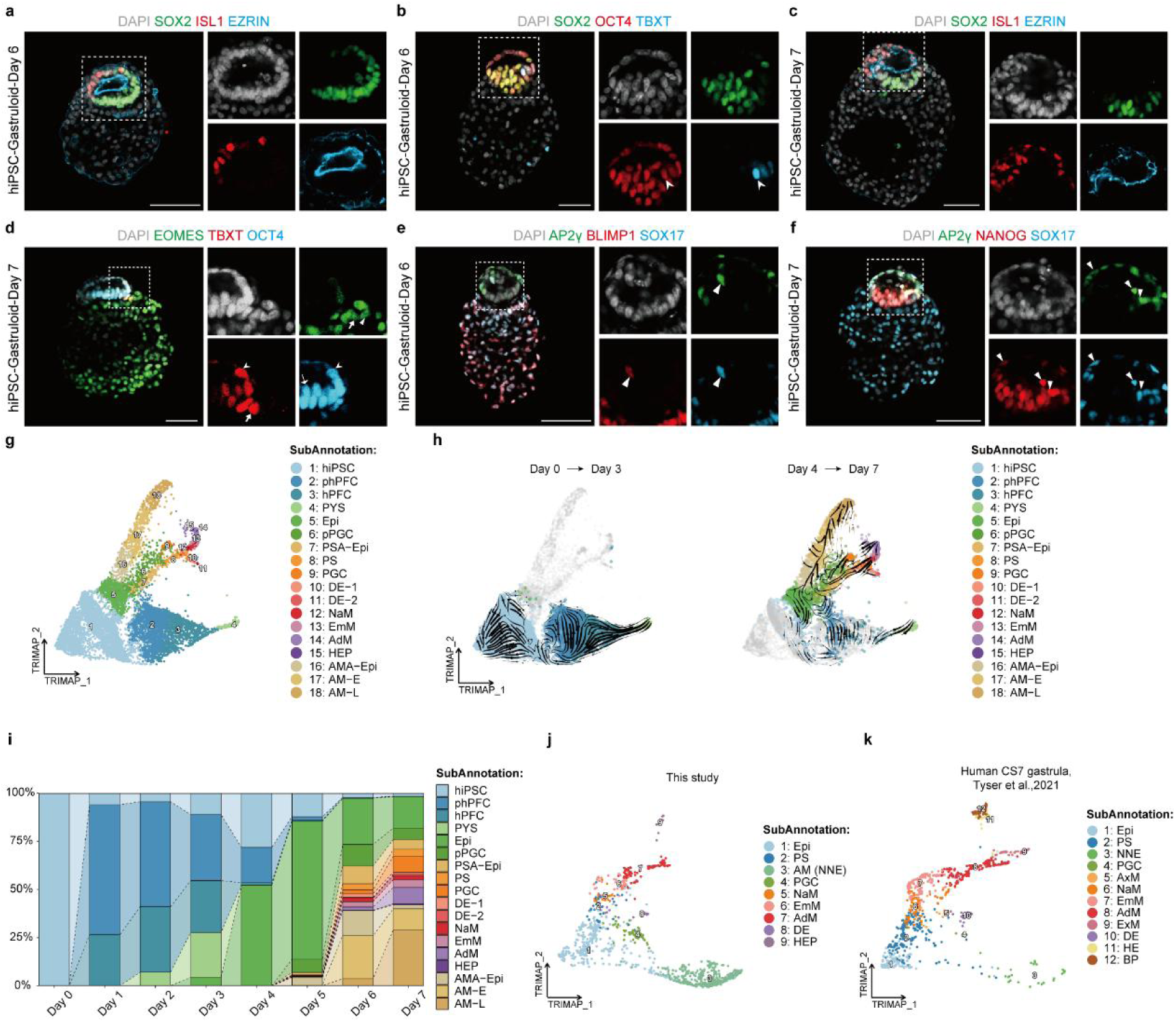
Fate determination of epiblast development. (a) IF staining for epiblast marker SOX2 (green), amnion cell marker ISL1 (red) and amnion cavity marker EZRIN (blue) of the human gastruloid (based on hiPSC) on day 6. Nuclei were counterstained with DAPI (grey). Scale bar, 100 μm. (b) IF staining for epiblast marker SOX2 (green), OCT4 (red) and primitive streak marker TBXT (blue) of the human gastruloid (based on hiPSC) on day 6. Nuclei were counterstained with DAPI (grey). White arrowhead, OCT4^+^TBXT^+^ primitive streak. Scale bar, 100 μm. (c) IF staining for epiblast marker SOX2 (green), squamous amniotic epithelium marker ISL1 (red) and amnion cavity marker EZRIN (blue) of the human gastruloid (based on hiPSC) on day 7. Nuclei were counterstained with DAPI (grey). Scale bar, 100 μm. (d) IF staining for epiblast marker OCT4 (blue), primitive streak marker TBXT (red) and endoderm marker EOMES (green) of the human gastruloid (based on hiPSC) on day 7. Nuclei were counterstained with DAPI (grey). White solid arrow, only OCT4^+^ epiblast. White arrowhead, OCT4^+^TBXT^+^ primitive streak. White arrow, TBXT^+^EOMES^+^ cell. White triangle arrow, only EOMES^+^ definitive endoderm. Scale bar, 100 μm. (e, f) IF staining for the primordial germ cell marker SOX17 (blue), AP2γ (green) co-stained with BLIMP1 (red) of the human gastruloid (based on hiPSC) on day 6 and NANOG (red) of the human gastruloid (based on hiPSC) on day 7. Nuclei were counterstained with DAPI (grey). White triangle arrow, primordial germ cell. Scale bars, 100 μm. (g) TriMap revealed 18 subtypes. hiPSC, human induced pluripotent stem cell; hPFC, pluripotent founder cell; phPFC, pre-hPFC; PYS, primary yolk sac; Epi, epiblast; PSA-Epi, primitive streak anlage-epiblast; PS, primitive streak; PGC, primordial germ cell; pPGC, precursor of PGC; DE, definitive endoderm; NaM, nascent mesoderm; EmM, emergent mesoderm; AdM, advanced mesoderm; HEP, haemato-endothelial progenitor; AMA-Epi, amnion anlage-epiblast; AM-E, amnion-Early; AM-L, amnion-Late. (h) RNA velocity analysis inferring the development of epiblast. (i) Bar plot showing the proportions of 18 subtypes in each sampling day. (j-k) TriMaps of integration of Epi derivatives in gastruloids on day 7 (j) and human CS7 gastrula dataset^13^ (k). NNE, non-neural ectoderm; AxM, axial mesoderm; HE, hemogenic endothelium; BP, blood Progenitors.

To track the differentiation trajectory of epiblast in our gastruloids, we calculated the DDGs of epiblast along the trajectories (Extended Data Fig. 12a-c, Supplementary Table 2). We found that C3 DDGs were crucial for the development of epiblast into PGC. C4 DDGs were highly associated with gastrulation development, and C5 DDGs were up regulated during the formation of the amnion. Interestingly, C2 DDGs were highly expressed during the developmental processes of PGC, PS, and amnion. Furthermore, pPGC, PSA-Epi, and AMA-Epi exhibited a high similarity in certain transcriptomic features (Extended Data Fig. 10d). Together, these results suggested that precursor cells of these cell types underwent a transient phase of similar transcriptome states, which was consistent with previous reports^28–30^. Moreover, our results indicated that the PGC originated from the epiblast at the caudal end of the embryonic disc. To further explore how the lineage of PGC specification was established, we performed PAGA analysis. We found that pPGC highly correlated with both AMA-Epi and PSA-Epi in developmental relationships (Extended Data Fig. 12b). We next calculated the DDGs of gastrulation along the trajectories (Extended Data Fig. 12d-f, Supplementary Table 2). We found that C2 DDGs were highly correlated with the development of mesoderm, while C3 DDGs were highly expressed when DE developed.

Together, these findings clearly demonstrated that the morphology and molecular properties of epiblast in our gastruloid determined how it differentiated into the three germ layers and germ cells.

### Research models for disorders of embryonic development

To evaluate whether our gastruloids are suitable to study the effects of drugs on embryonic development, we investigated the effects of thalidomide (THD) incubation on gastrula development in our model. THD is a teratogenic drug that was once widely used because of its strong efficacy in the treatment of vomiting during pregnancy^31^. The teratogenicity of THD displays a complex interplay with the species in question, highlighting observed discrepancies between human and other animal models^32–37^. Here, we designed a toxicity test for our gastruloids to investigate the impact of THD exposure on early embryonic development.

The plasma concentration in human taking THD is usually between 1-6 μM^38–40^. We designed normal group, DMSO-control group, 5 μM-THD group, and 10 μM-THD group for experiments. We started to add THD from day 0 during the gastruloids induction process. We found that on day 7, the aggregates treated with 5 μM and 10 μM THD failed to form the amniotic cavity and yolk sac structure (Fig. 4a). We also observed that these aggregates were significantly smaller in volume than those of normal and DMSO-control groups (Fig. 4a). To track cell proliferation and volume in the aggregates over time, we performed statistical analysis for seven consecutive days. We found that with increasing drug concentrations, the cell number and volume of aggregates decreased (Fig. 4b). By performing nuclear staining of gastruloids, we found that the cells in THD-treated groups were arranged in a disorderly and compact manner (Fig. 4a). Our IF staining in 5 μM-THD group showed that the epiblast failed to undergo ordered polarization and formed negligible amounts of amnion, presumably because the epiblast failed to transform into a squamous epithelial-like morphology (Extended Data Fig. 13a, e). On day 7, the majority of aggregates exhibited an absence of TBXT^+^ gastrulating cells (Extended Data Fig. 13b, e). Only in a small number of aggregates, a subset of cells underwent gastrulation and were found dispersed within the epiblast (Extended Data Fig. 13e). In addition, we found that the hypoblast development was chaotic (Extended Data Fig. 13c, d, f, g). We found that OTX2 was widely expressed in the hypoblast of 5 μM-THD group, while it was locally expressed with polarity localization in the hypoblast of normal group (Extended Data Fig. 13c, f). Furthermore, we found that the GATA6^+^ endoderm enveloped the LAMC1^+^ mesoderm in 5 μM-THD group, exhibiting an abnormal yolk sac (Extended Data Fig. 13d, g).

**Fig. 4.**
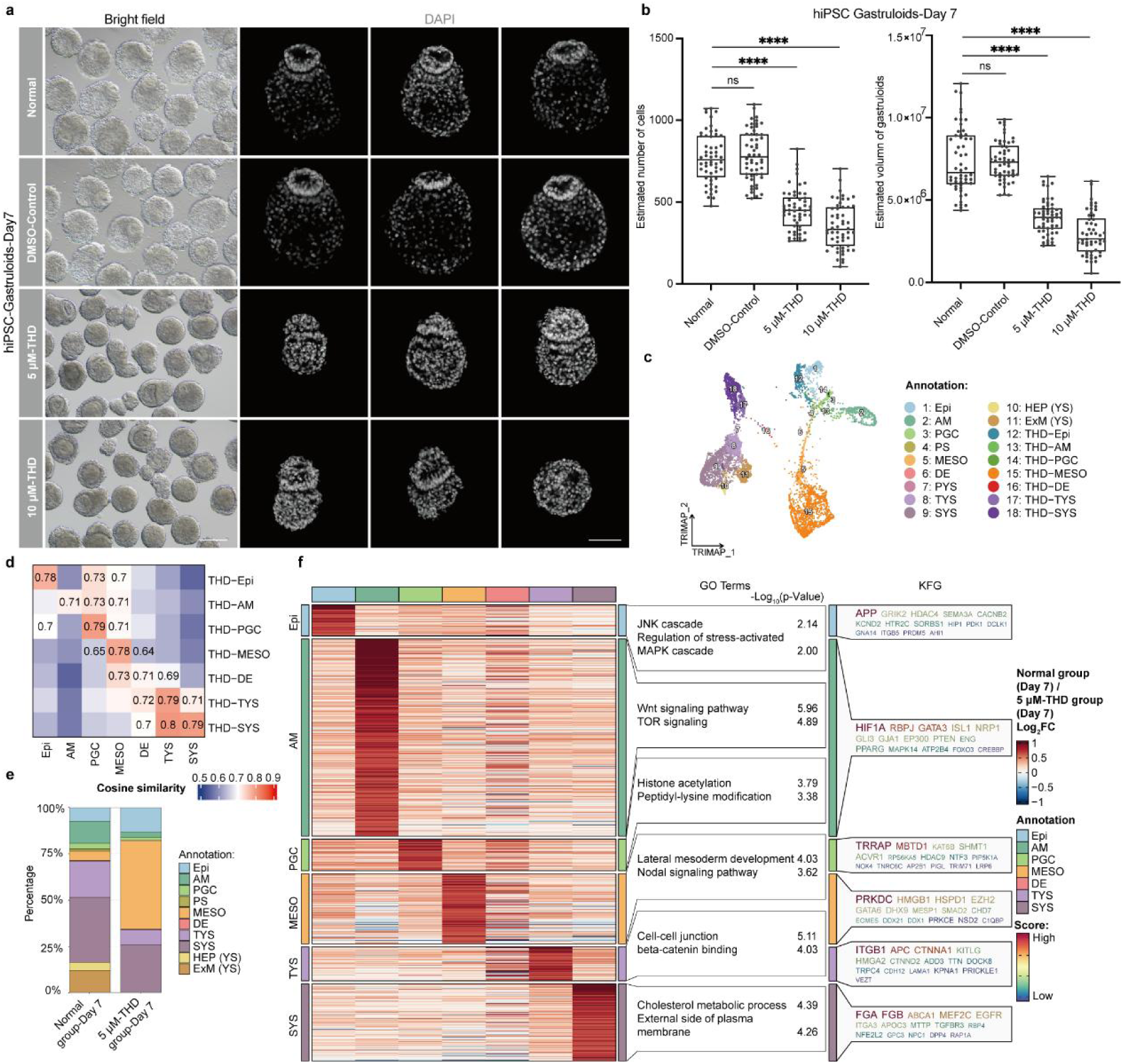
The effect of Thalidomide on gastruloids development. (a) Brightfield images (left) and DAPI staining (right) for human gastruloids (based on hiPSC) in four groups on day 7. Nuclei were counterstained with DAPI (grey). Scale bars, 200 μm (left) or 100 μm (right). (b, c) Quantification of total cell numbers (left) and volume (right) in per gastruloid (based on hiPSC) in four groups on day 7; ****P < 0.0001(Welch two-sided (left), Unpaired two-sided (right), two-sample t-test). Centre line, mean; error bar, S.D; n= 54. (c) TriMap showing cell types of gastruloids with 5 μM THD treatment integrated with the normal group on day 7. Epi, epiblast; AM, amnion; PS, primitive streak; PGC, primordial germ cell, MESO, mesoderm; DE, definitive endoderm; PYS, primary yolk sac; TYS, transitional yolk sac; SYS, secondary yolk sac; HEP, haemato-endothelial progenitor; ExM, extraembryonic mesoderm. (d) Heatmap showing the cosine similarity between cell types of the same categories with and without 5 μM THD treatment. (e) Bar plot showing the proportion of cell types on day 7 with and without 5 μM THD treatment. (f) Heatmap of the average expression pattern of DEGs disturbed in each cluster that differently affected by 5 μM THD treatment (left panel). The related Gene Ontology (GO) enrichment terms (middle panel) and key functional genes (KFGs) (right panel) are shown.

To better understand how THD affects gastruloid development at the molecular level, we performed scRNA-Seq on aggregates of 5 μM-THD group on day 7, with a total of over 3000 cells passing our stringent quality control. We integrated these data with that of our normal group (Fig. 4c, Extended Data Fig. 14a). Despite the fact that 5 μM-THD aggregates were similar to normal gastruloids in cell types (Fig. 4d, Extended Data Fig. 14b, Supplementary Table 1), we found that their proportion and transcript levels of cells differed significantly (Fig. 4e, f, Extended Data Fig. 14c, Supplementary Table 1). When we next analyzed the proportion of cells in 5 μM-THD aggregates, we found that these aggregates preferentially developed into mesoderm. Mesoderm cells accounted for approximately 50% of the total cell numbers, whereas the development of amnion decreased, with amniotic cells accounting for less than 3% (Fig. 4e). When we analyzed DEGs between the normal group and the 5 μM-THD group conserved in cell types, we found that biological terms related to mitochondria and ribosomes enriched in 5 μM-THD group (Extended Data Fig. 14c, Supplementary Table 1). We also found that the normal group enriched in biological terms related to amnion development, such as regulation of cell-matrix adhesion and cell junction assembly, providing further evidence of a noticeable impact on amnion development following THD treatment (Extended Data Fig. 14c, Supplementary Table 1). When we analyzed unique DEGs of cell types between the normal group and the 5 μM-THD group, we found that the amnion exhibited the highest sensitivity to the THD, followed by the SYS and the MESO (Fig. 4f, Supplementary Table 1). Furthermore, we found that the unique DEGs of clusters were enriched in many signal pathways related to early embryo development, such as the MAPK, Wnt, and Nodal signaling pathways (Fig. 4f, Supplementary Table 1). Lastly, to assess the effect of THD on gastruloids development, we focused on differences in transcriptome of the MAPK and Wnt pathways. When we performed gene set enrichment analysis (GSEA), we found that the MAPK and Wnt signaling pathways were significantly downregulated in 5 μM-THD group (Extended Data Fig. 15a, b). Furthermore, compared to the normal group, 5 μM-THD group displayed prominent downregulation of numerous genes linked with these signaling pathways (Extended Data Fig. 15c, d). Our results were consistent with those of a previous study using chicken embryos, supporting the results that inhibition of the Wnt pathway is responsible for limb defects following THD administration^41^.

Together, these findings strongly indicated that THD-treatment results in disordered development of gastruloids.

## Discussion

Here, we report a novel gastruloid that mimics the key events occurring during human gastrula, such as hypoblast specialization, anterior visceral endoderm polarization, amniotic cell epithelialization, extraembryonic mesoderm formation, primordial germ cell specification, and gastrulation from CS4 to early CS7.

Human embryogenesis involves two phases of yolk sac development: a transient primary yolk sac derived from the hypoblast, followed by the rapid replacement with a secondary yolk sac during gastrulation^18^. However, the mechanism underlying this replacement is still unknown^5^. Here, we found that PYS-VE of gastruloids developed into pSYS by activating TGF-β and Wnt signalling pathways, and eventually differentiated into SYS-VE, SYS-PE, and SYS-ExM. Moreover, the SYS-PE was derived from the SYS-VE. These results suggest that pSYS constitute a prerequisite for the orderly development of the functional yolk sac in our gastruloids.

The ExM specialization prior to the formation of PS is characteristic of embryogenesis in primates, but the origin of the ExM remain unclear^18^. One hypothesis proposes that the hypoblast-derived PYS serves as a source for the ExM, which is supplemented by mesoderm from the gastrula^18,42^. Our gastruloids show that the PYS-ExM is derived from PYS-PE. Furthermore, PYS-ExM and pSYS differentiate into SYS-ExM through tExM. However, the molecular characteristics of PYS-ExM and SYS-ExM differ considerably. PYS-ExM is mainly similar to the transcriptome features of PYS-PE, while SYS-ExM was similar to SYS-PE.

Experiments on placental mammals have consistently shown that the hypoblast gives rise to both the ExM and the HEP^26,43^. However, it remained unclear whether the HEP arises directly from the hypoblast or is formed through the ExM derived from the hypoblast^44^. Our research suggests that at the molecular level, the SYS-PE derived from PYS-VE is responsible for the earliest hematopoiesis during hypoblast differentiation in gastruloids.

Various explanations have currently been put forward that human amnion segregates prior to PGC formation or a “dual origin” elucidates PGCs from both the amnion and posterior epiblast^7,45–48^. The gastruloids we established shows that PGC was derived from the epiblast. This result supports the notion that the PGC originates from the posterior epiblast of the embryonic disc.

Preclinical studies, which test developmental and reproductive toxicity, are performed on lab animals to anticipate and mitigate any negative impacts drugs may have on human reproduction and growth. Nonetheless, these studies necessitate the use of a significant quantity of animals^4^. Furthermore, certain drugs may demonstrate intricate interactions with the respective species. Extensive toxicological evaluations reveal that the use of THD by expectant mothers is associated with severe congenital malformations in offspring, such as thumb loss, limb dysplasia, neurodevelopmental retardation, and congenital heart disease^31^. Several stem cell-based models have previously been utilized to investigate the effects of THD^37,49–51^; However, most of these models lack the crucial spatiotemporal and morphological context of the developing embryo. Furthermore, the precise effects of THD on early embryonic development of human remains unclear. Here, we found that the teratogen THD has a dose-dependent effect on the dysplasia of gastruloids. As this model simulates key morphological and molecular information during gastrulation, it makes reproductive toxicology testing of drugs more sensitive and accurate. In particular, our model avoids the potential legal, regulatory, and ethical concerns, as it lacks the potential to form a whole organism.

Although our gastruloid mimics a majority of cell types present in the human CS7 gastrula, it falls short in replicating the axial mesoderm cell type. Furthermore, our gastruloid is non-integrative and does not contain trophoblast cells, making it unsuitable for the screening of drugs that lead to abnormal trophoblast development.

In conclusion, the sequential induction process we established, derived from human pluripotent stem cell, faithfully recapitulates the early stages of human embryonic development, distinguishing it from other assembly models involving multiple cell combinations. Our model enables a detailed characterization of the molecular trajectory origins of differentiating cells, allowing for a more precise delineation of cellular fate determination paradigms. Our research convincingly demonstrates that our model is well-suited to accurately predict the potential interference of newly developed drugs with human embryonic development.

## Acknowledgements

We thank Xi Wang from Nanjing Medical University for his valuable discussions and comments. We thank Carl Zeiss AG (Asia Pacific) for providing us with imaging and processing services for the light sheet images.

## Author contributions

J.S. initiated the project. J.S. and Y.Y. organized and supervised the entire project. J.S., Y.Y., Y.G. and J.C. wrote the manuscript. Y.G. designed the experiments. Y.G., M.C., and P.Z. performed the experiments. Y.C. and Z.L. constructed single-cell transcriptome library. J.C. and H.Z. analysed RNA-seq data.

## Data and materials availability

All data generated or analyzed during this study have been uploaded to the National Center for Biotechnology Information (NCBI) databases : https://dataview.ncbi.nlm.nih.gov/object/PRJNA956322. Access to the data is free and open after release.

## Supplementary Information

Materials and Methods

Tables 1-3

Movies 1 to 3

References 1-14

---

**Extended Data Fig. 1.**
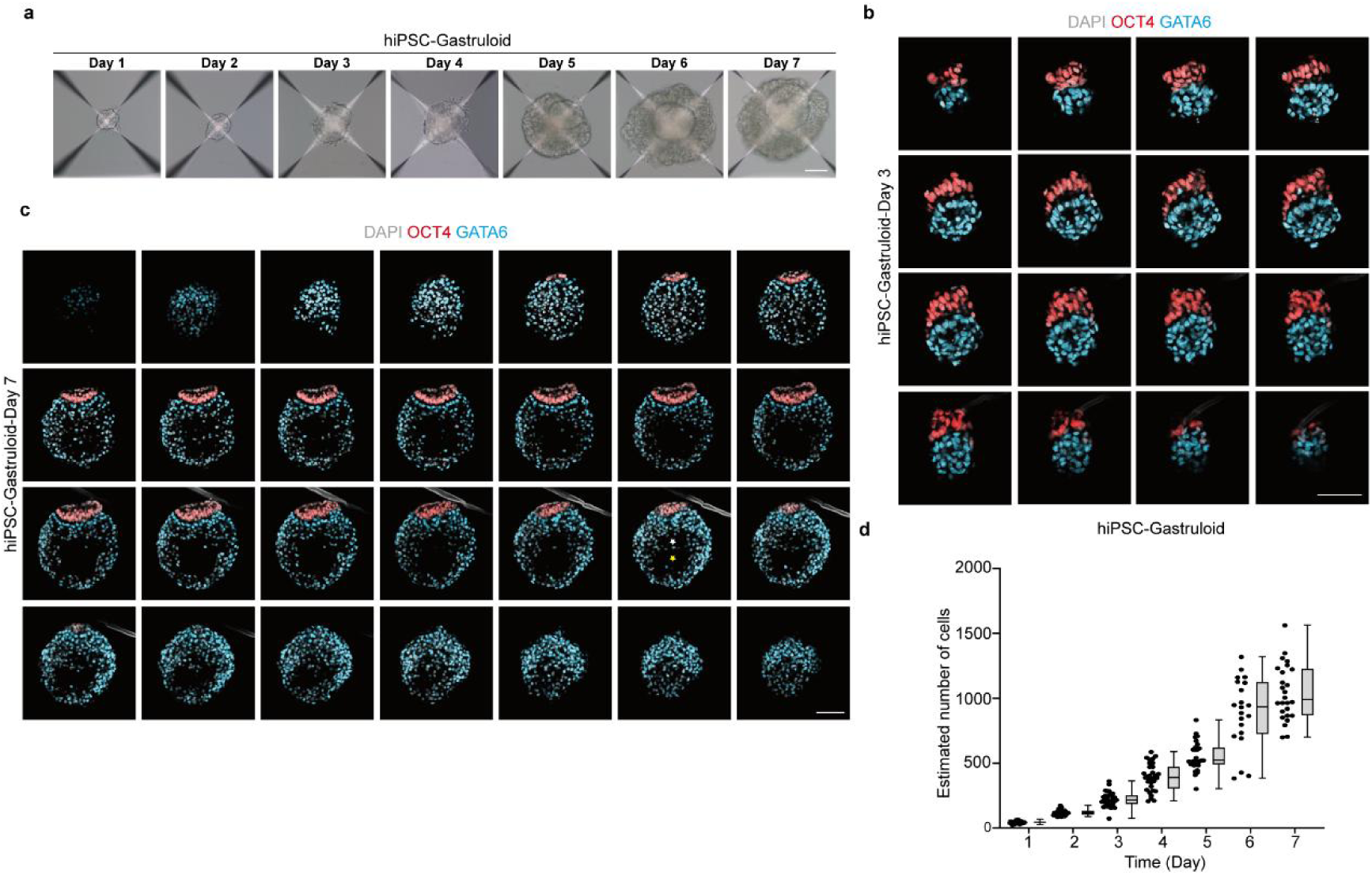
Morphological characterization of human gastruloids. (a) Brightfield images of the human gastruloid (based on hiPSC) from day 1 to 7. Scale bar, 100 μm. (b) Series of confocal z-sections of the human gastruloid on day 3 (based on hiPSC) stained for epiblast marker OCT4 (red) and hypoblast marker GATA6 (blue). Nuclei were counterstained with DAPI (grey). Scale bar, 100 μm. (c) Series of confocal z-sections of the day 7 human gastruloid (based on hiPSC) stained for epiblast marker OCT4 (red) and hypoblast marker GATA6 (blue). Nuclei were counterstained with DAPI (grey). Yellow asterisk, primary yolk sac; White asterisk, secondary yolk sac. Scale bar, 100 μm. (d) Quantification of total cell numbers in per developing human gastruloid (based on hiPSC) from day 1 to 7. Error bar, Mean with range; box, interquartile range; n= 29,27,36,32,28,21,26 human gastruloids from day 1 to 7.

**Extended Data Fig. 2.**
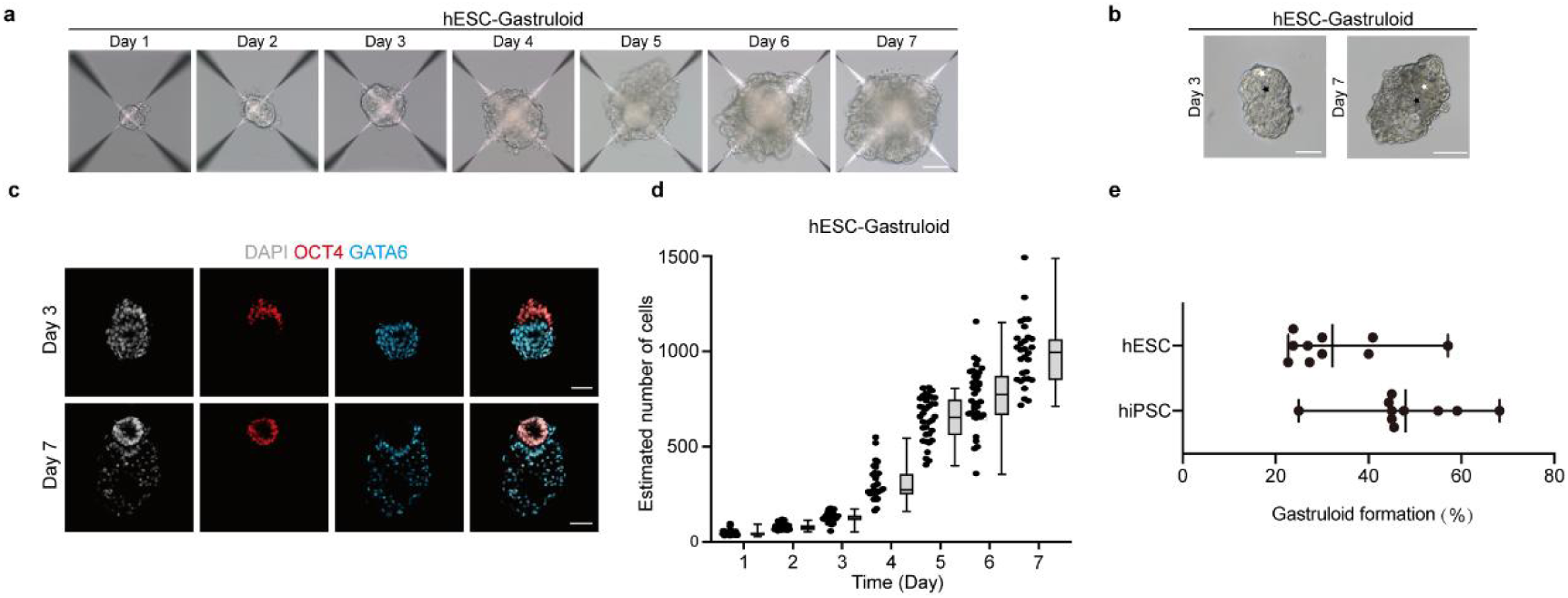
Morphological characterization of hESC-based gastruloids. (a) Brightfield images of representative the human gastruloid from day 1 to 7. Scale bar, 100 μm. (b) Brightfield images of representative the human gastruloid on day 3 and 7. White asterisks, epiblast. Black asterisks, hypoblast. Scale bars, 100 μm. (c) IF staining for epiblast marker OCT4 (red) and hypoblast marker GATA6 (blue) of the human gastruloid on day 3 and 7. Nuclei were counterstained with DAPI (grey). Scale bars, 50 μm (d) Quantification of total cell numbers in per developing human gastruloid from day 1 to 7. Error bar, Mean with range; box, interquartile range; n= 36,33,37,31,35,37,30 human gastruloids from day 1 to 7. (e) Quantification of the percentage of gastruloids including epithelial polarization, gastrulation and secondary yolk sac cavity formation for hiPSC and hESC induced by two-stage protocol. Error bar: Mean with range, n=10 per cell line.

**Extended Data Fig. 3.**
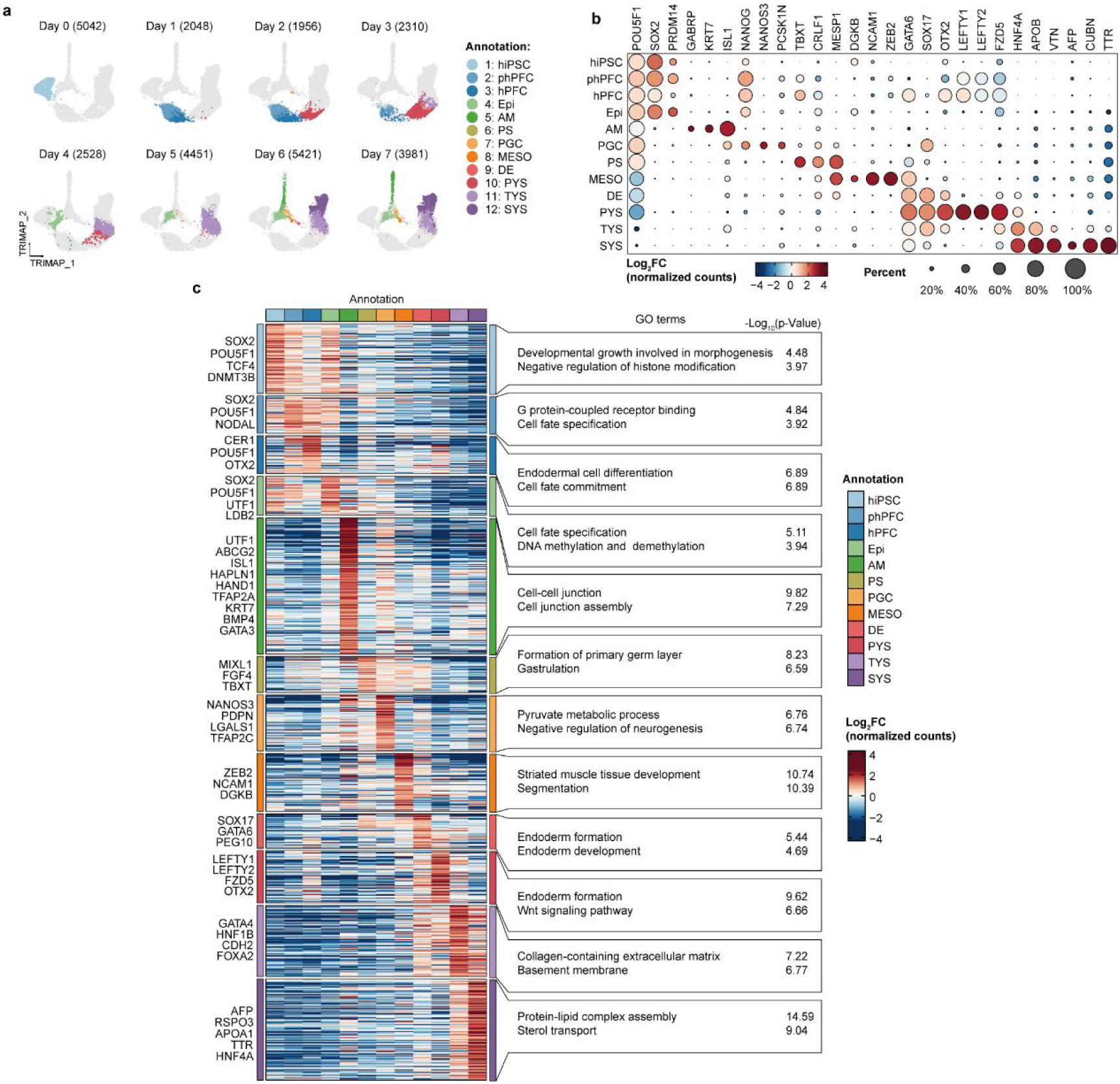
Transcriptome features and clustering criteria of gastruoids. (a) TriMap showing the proportions of 12 major cell types in each sampling day. (b) Bubble plot representing the frequency of expression and average expression of key lineage-associated genes in identified clusters in Fig.1d. (c) Heatmap of the average expression pattern of DEGs of each cell type (middle panel). The representative marker genes (left panel) and related GO enrichment terms (right panel) were shown. hiPSC, human induced pluripotent stem cell; hPFC, human pluripotent founder cell; phPFC, pre-hPFC; Epi, epiblast; AM, amnion; PS, primitive streak; PGC, primordial germ cell, MESO, mesoderm; DE, definitive endoderm; PYS, primary yolk sac; TYS, transitional yolk sac; SYS, secondary yolk sac.

**Extended Data Fig. 4.**
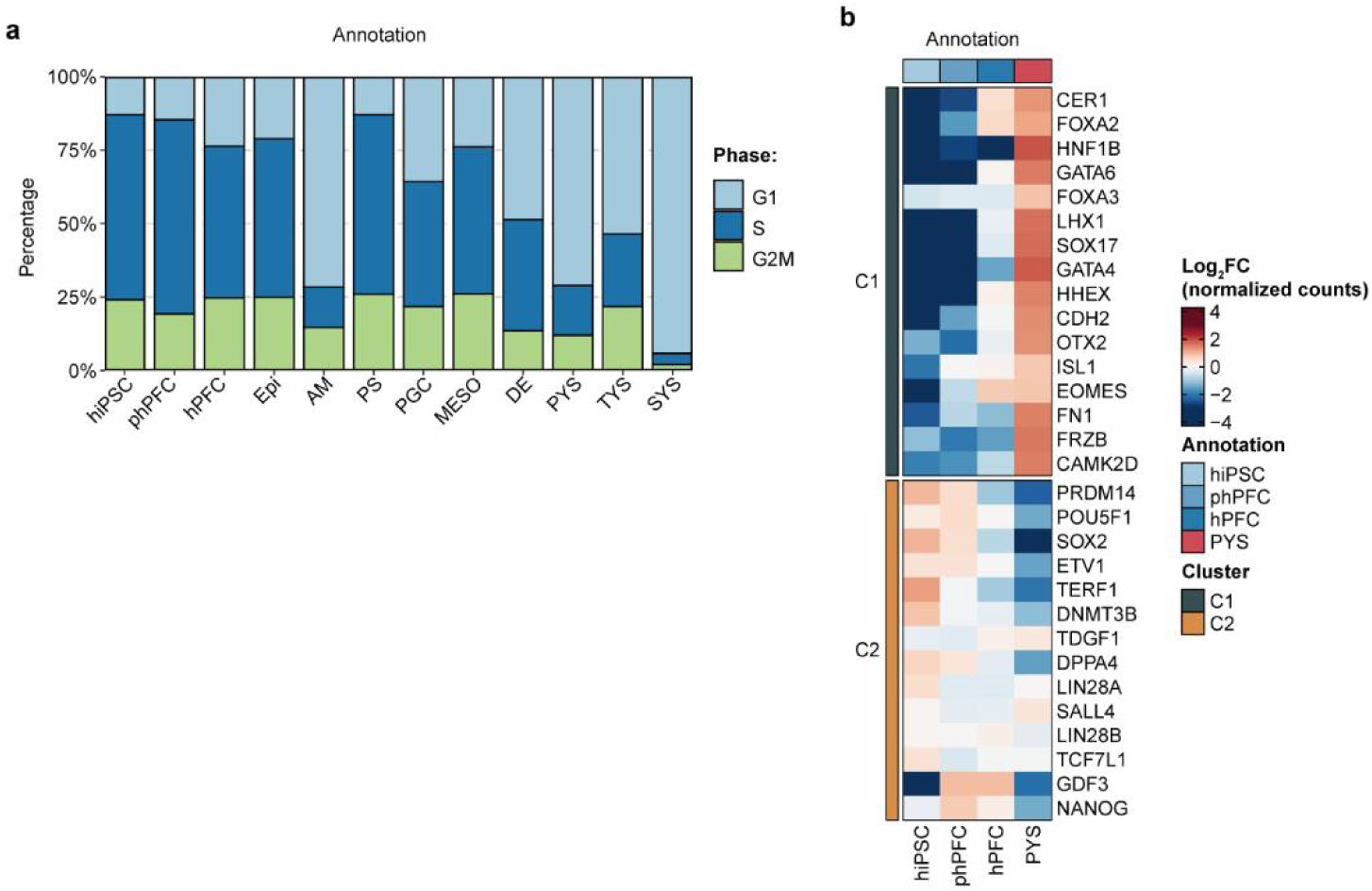
Transcriptome features of gastruoids. (a) Bar plot representing the percentage of cells in G1, S, or G2M phase of each major cell type. (b) Heatmap showing the different expression patterns of notable genes during generation of hPFC from Nakanishi et al.^10^. C1 marked genes related to endoderm formation, and C2 marked genes related to pluripotency.

**Extended Data Fig. 5.**
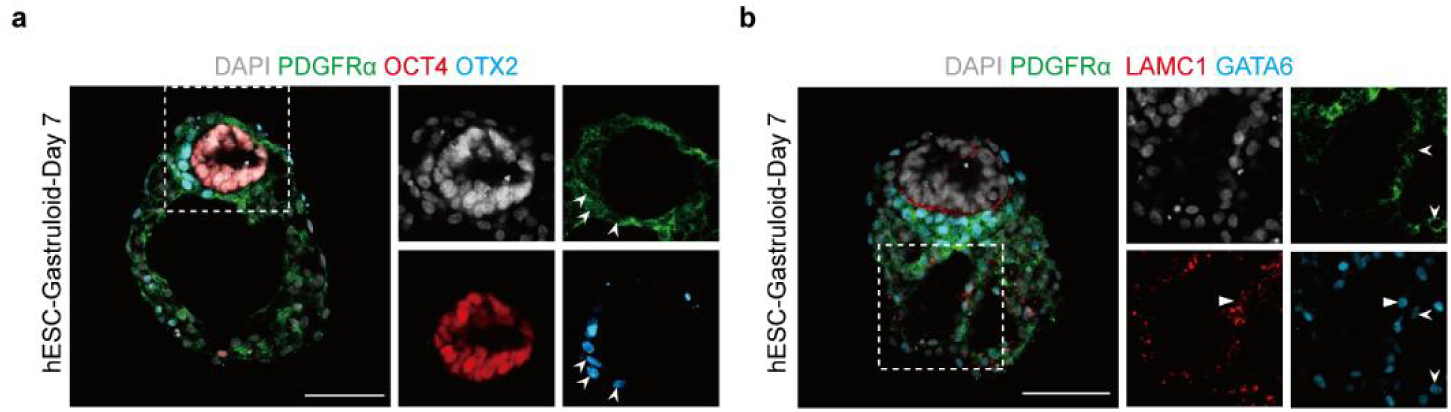
Morphological characterization of the hypoblast development in hESC-based gastruloids. (a) IF staining for epiblast marker (red) OCT4, hypoblast/anterior visceral endoderm marker (blue) OTX2 and secondary yolk sac marker (green) PDGFRα of the human gastruloid (based on hESC) on day 7. Nuclei were counterstained with DAPI (grey). White arrowhead, anterior visceral endoderm. Scale bar, 100 μm. (b) IF staining for hypoblast marker (blue) GATA6, secondary yolk sac marker (green) PDGFRα and extraembryonic mesoderm marker (red) LAMC1 of the human gastruloid (based on hESC) on day 7. Nuclei were counterstained with DAPI (grey). White arrowhead, secondary yolk sac. White triangle arrow, extraembryonic mesoderm. Scale bar, 100 μm.

**Extended Data Fig. 6.**
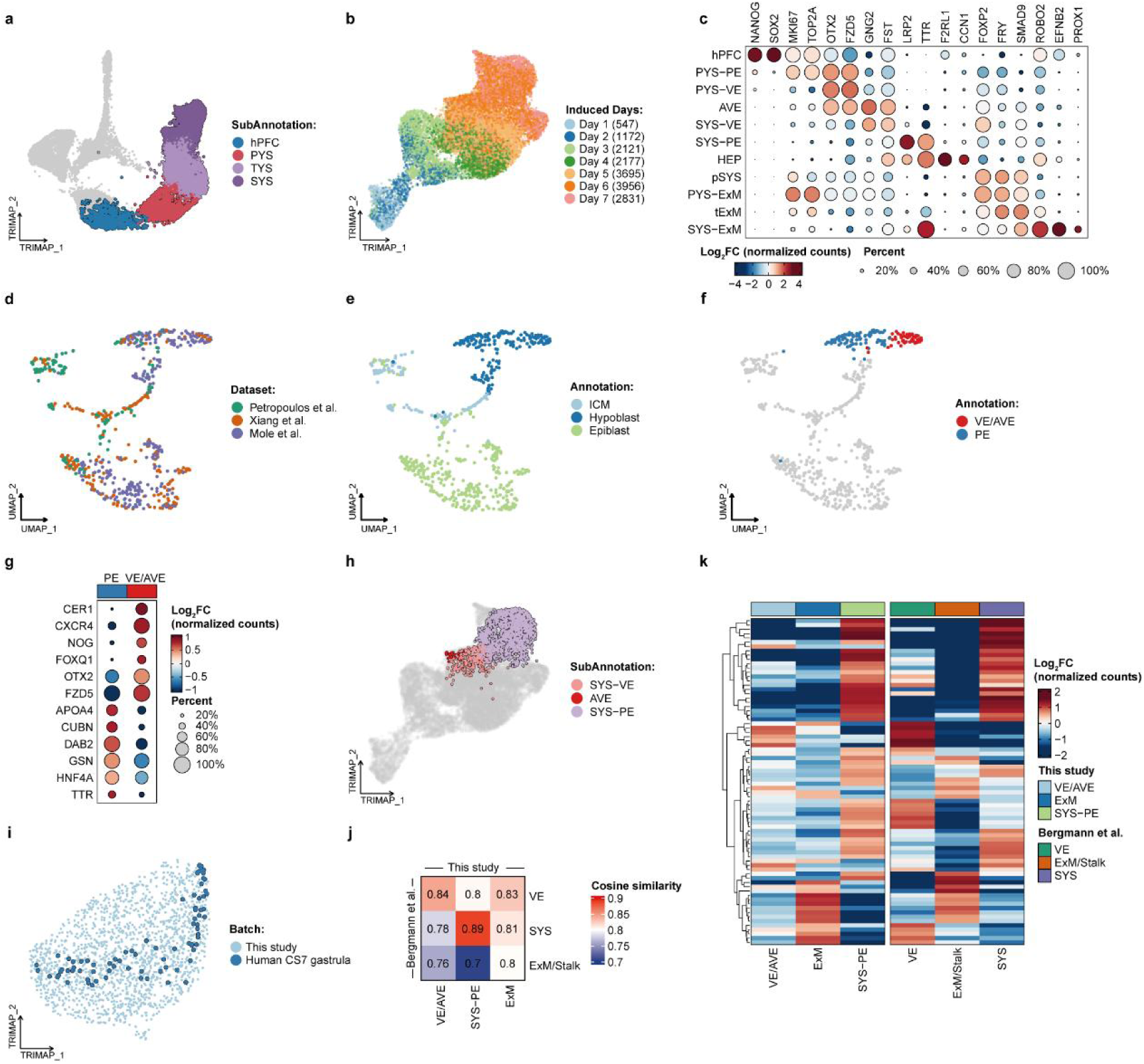
Clustering criteria and similarity analysis of derivatives of epiblast in gastruloids. (a) TriMap showing distribution of hypoblast derivatives in Fig. 1d with a highlighted border. (b) TriMap showing the distribution of hypoblast derivatives at sampling days. (c) Bubble plot representing the frequency of expression and average expression of key lineage-associated genes in identified clusters in Fig.2f. (d) UMAP presenting three human embryo reference datasets consisting of datasets for human pre-implantation embryos^20^, IVC human embryos^19^, and IVC human post-implantation embryos^15^. (e) UMAP showing the cell annotations retrieved from the query reference. (f) UMAP showing the VE/AVE and PE parts present in the integrated human embryonic reference distinguished according to the representative gene in (g). (g) Bubble plot representing the frequency of expression and average expression of specific marker genes of VE and PE. (h) TriMap showing the distribution of cells integrated with the human CS7 gastrula dataset^13^ with highlighted borders in Fig. 2f. (i) TriMap showing the distribution of human CS7 gastrula dataset^13^ and data of this study. (j) Heatmap showing the cosine similarity of VE, SYS and ExM/Stalk in marmoset datasets versus VE/AVE, SYS-PE and ExM in gastruloids, calculated from shared DEGs. (k) Heatmap of the average expression of shared DEGs used in (j).

**Extended Data Fig. 7.**
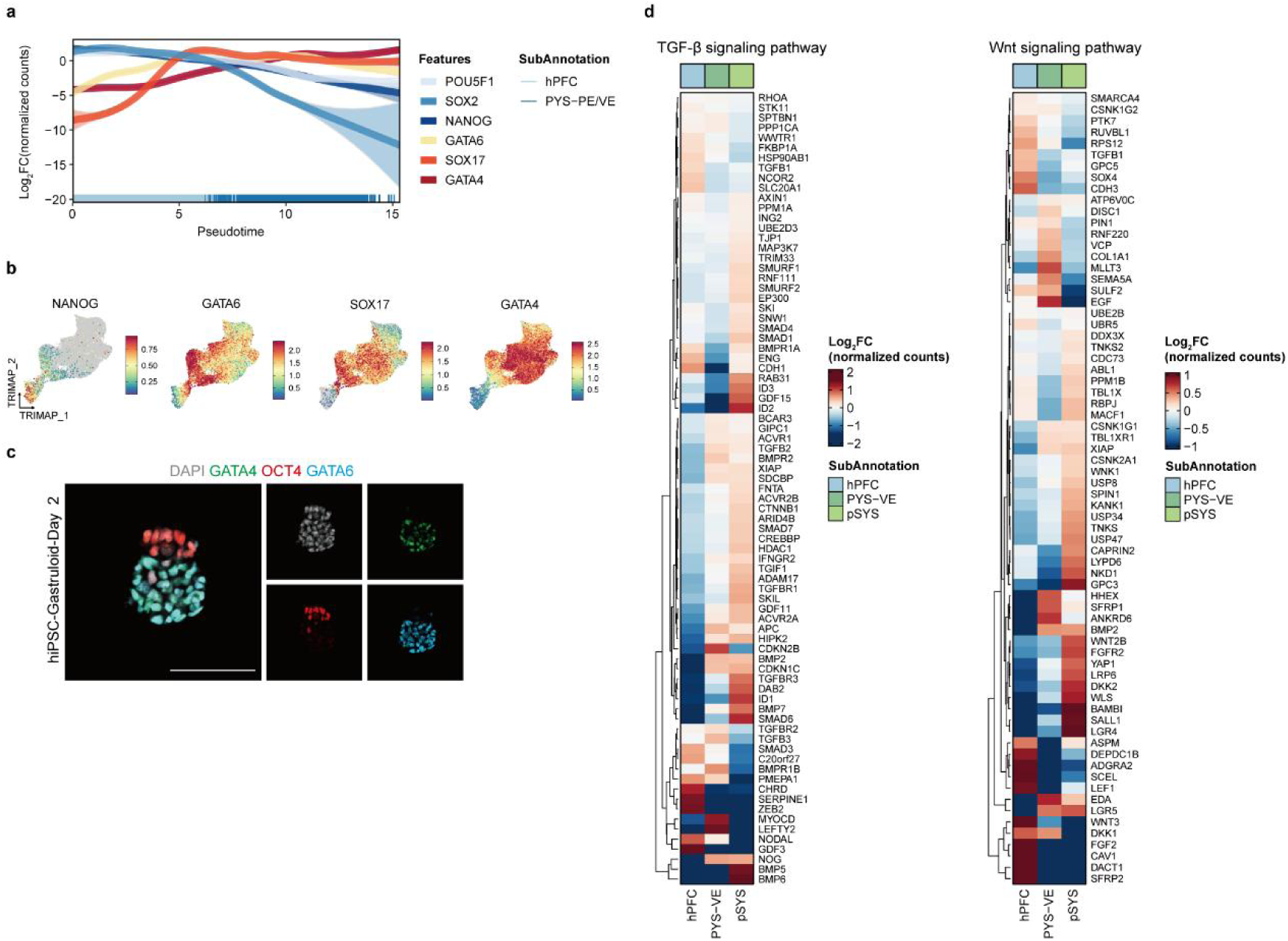
Transcriptome features of the hypoblast differentation. (a) Line plot showing expression of genes related to pluripotency and endoderm formation during hPFC development to PYS. (b) TriMap showing expression of pluripotent marker NANOG and sequential upregulation of hypoblast specifiers GATA6, SOX17 and GATA4. (c) IF staining for epiblast marker OCT4 (red), hypoblast marker GATA6 (blue) and GATA4 (green) of the human gastruloid (based on hiPSC) on day 2. Nuclei were counterstained with DAPI (grey). Scale bar, 100 μm. (d) Heatmaps showing expression of genes in TGF-β (ref: GO:0030511) (left) and WNT (ref: GO:0030177) (right) signaling pathways in indicated cell types.

**Extended Data Fig. 8.**
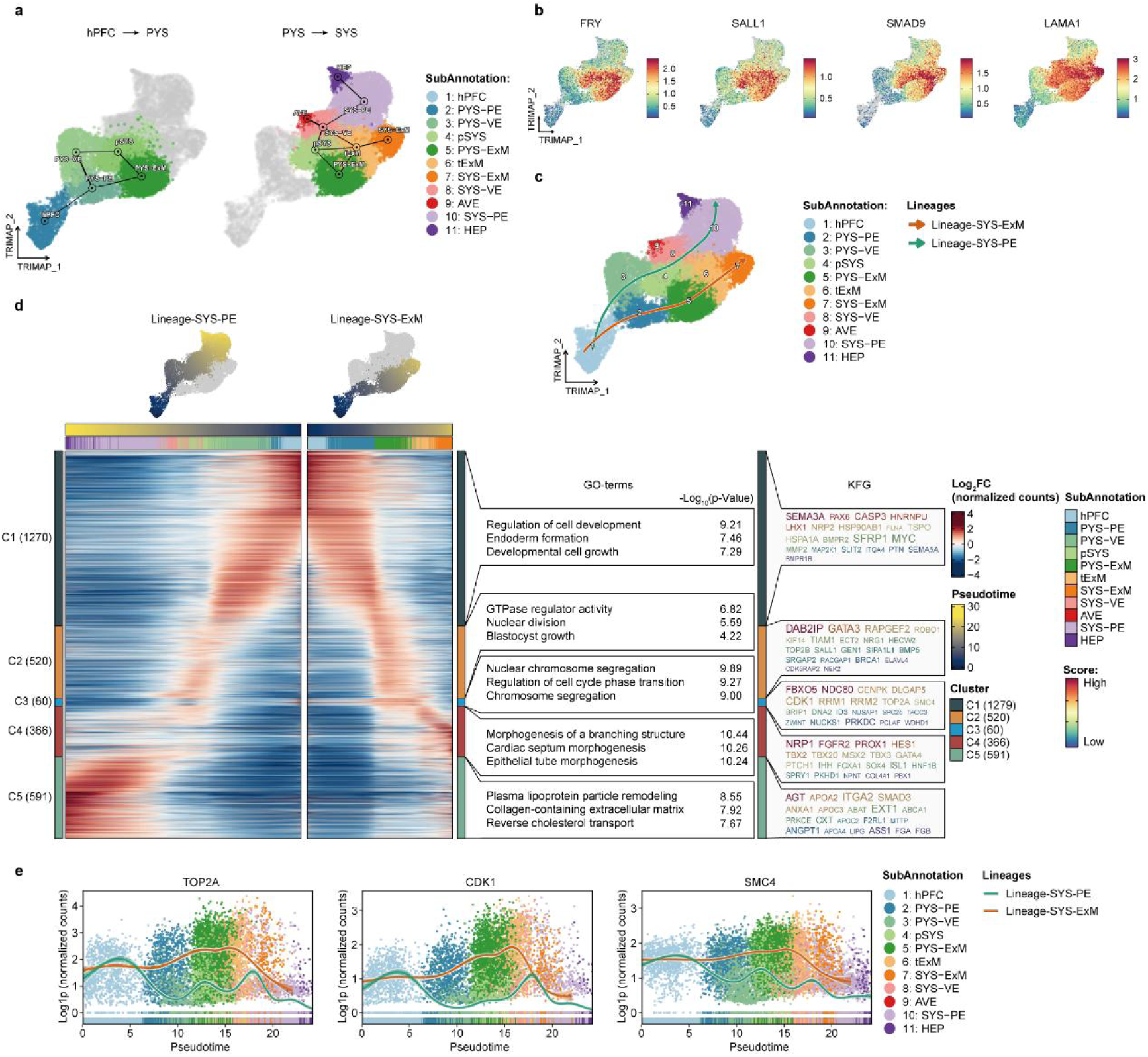
scRNA-seq analysis of the hypoblast development trajectories. (a) Divided into hPFC to PYS and PYS to SYS for PAGA analysis and overlaid on TriMaps. The threshold for connection of cell types was set to 0.05 (left) and 0.23 (right). (b) TriMaps showing expression of mesoderm-related genes. (c) TriMap showing cell trajectories inferred by Slingshot where cells underwent transitions from hPFC to HEP and to SYS-ExM. (d) Heatmap showing expression pattern dynamics (left panel) alongside the developmental trajectories from hPFC to HEP and to SYS-ExM with enriched pathways (middle panel) and KFGs (right panel). TriMaps colored by pseudotime were annotated above the corresponding trajectories. (e) Line plot showing the expression of genes related to cell proliferation on trajectories, and the points represent different subtypes.

**Extended Data Fig. 9.**
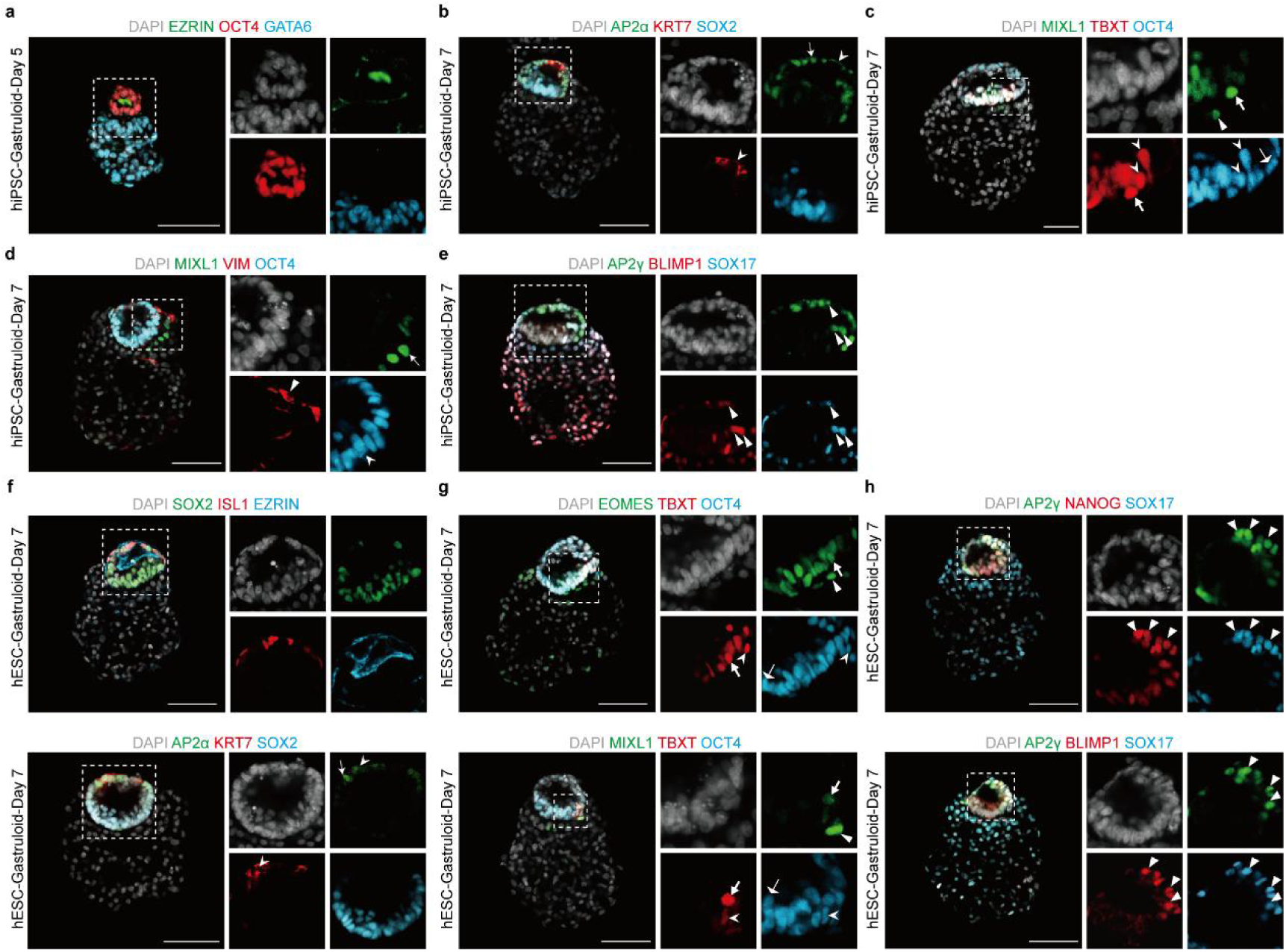
Morphological characterization of derivatives of the epiblast. (a) IF staining for epiblast marker OCT4 (red), hypoblast marker GATA6 (blue) and epithelial polarization marker EZRIN (green) of the human gastruloid (based on hiPSC) on day 5. Nuclei were counterstained with DAPI (grey). Scale bar, 100 μm. (b) IF staining for epiblast marker SOX2 (blue), amnion cell marker AP2α (green) and KRT7 (red) of the human gastruloid (based on hiPSC) on day 7. Nuclei were counterstained with DAPI (grey). White arrow, amnion cell. White arrowhead, squamous amniotic epithelium. Scale bar, 100 μm. (c) IF staining for epiblast marker OCT4 (blue), primitive streak marker TBXT (red) and mesoderm marker MIXL1 (green) of the human gastruloid (based on hiPSC) on day 7. Nuclei were counterstained with DAPI (grey). White solid arrow, only OCT4^+^ epiblast. White arrowhead, OCT4^+^ TBXT^+^ primitive streak. White arrow, TBXT^+^MIXL1^+^ cell. White triangle arrow, only MIXL1^+^ nascent mesoderm. Scale bar, 100 μm. (d) IF staining for epiblast marker (blue) OCT4, nascent mesoderm marker (green) MIXL1 and advanced mesoderm marker(red) VIM of the human gastruloid (based on hiPSC) on day 7. Nuclei were counterstained with DAPI (grey). White arrowhead, only OCT4^+^ epiblast. White arrow, MIXL1^+^ nascent mesoderm. White triangle arrow, VIM^+^ advanced mesoderm. Scale bar, 100 μm. (e) IF staining for primordial germ cell marker SOX17 (blue), AP2γ (green) and BLIMP1 (red) of the human gastruloid (based on hiPSC) on day 7. Nuclei were counterstained with DAPI (grey). White triangle arrow, primordial germ cells. Scale bar, 100 μm. (f) IF staining for epiblast marker SOX2 (green), amnion cell marker ISL1 (red), amnion cavity marker EZRIN (blue) (upper) and the epiblast marker SOX2 (blue), amnion cell marker AP2α (green) and KRT7 (red) (lower) of the human gastruloid (based on hESC) on day 7. Nuclei were counterstained with DAPI (grey). White arrow, amnion cell. White arrowhead, squamous amniotic epithelium. Scale bars, 100 μm. (g) Upper: IF staining for epiblast marker OCT4 (blue), primitive streak marker TBXT (red) and endoderm marker EOMES (green) of the human gastruloid (based on hESC) on day 7. Nuclei were counterstained with DAPI (grey). White solid arrow, only OCT4^+^ epiblast. White arrowhead, OCT4^+^TBXT^+^ primitive streak. White arrow, TBXT^+^EOMES^+^ cell. White triangle arrow, only EOMES^+^ definitive endoderm. Scale bar, 100 μm. Lower: IF staining for epiblast marker OCT4 (blue), primitive streak marker TBXT (red) and nascent mesoderm marker MIXL1 (green) of the human gastruloid (based on hESC) on day 7. Nuclei were counterstained with DAPI (grey). White solid arrow, only OCT4^+^ epiblast. White arrowhead, OCT4^+^TBXT^+^ primitive streak. White arrow, TBXT^+^MIXL1^+^ cell. White triangle arrow, only MIXL1^+^ nascent mesoderm. Scale bar, 100 μm. (h) IF staining for primordial germ cell marker SOX17 (blue), AP2γ (green) co-stained with NANOG (red) (upper) and BLIMP1 (red) (lower) of the human gastruloid (based on hESC) on day 7. Nuclei were counterstained with DAPI (grey). White triangle arrow, primordial germ cells. Scale bars, 100 μm.

**Extended Data Fig. 10.**
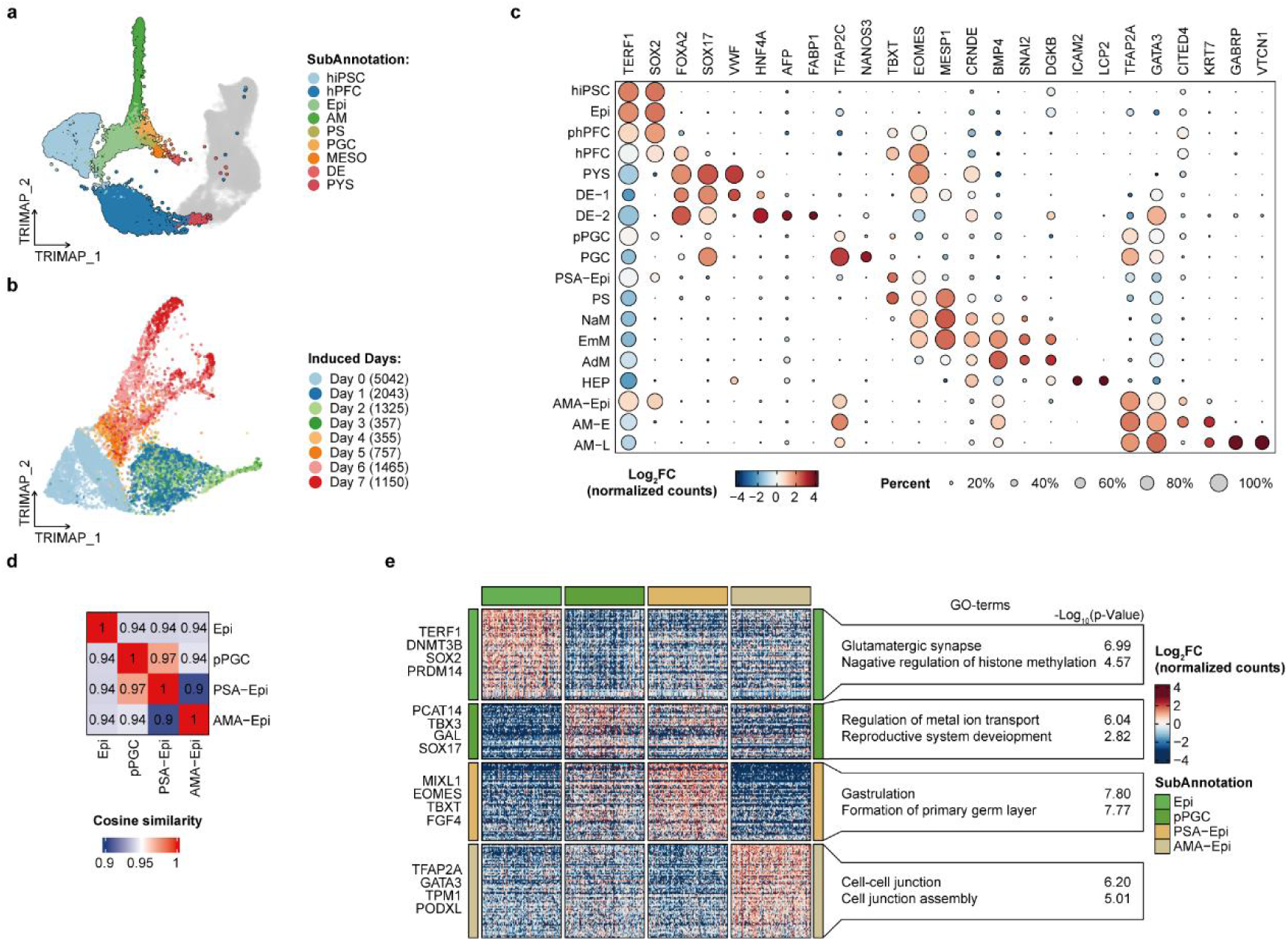
Transcriptome features and clustering criteria of the epiblast and its derivatives. (a) TriMap showing distribution of hiPSC, phPFC, hPFC, PYS and Epi derivatives in Fig. 1d with a highlighted border. (b) TriMap showing the distribution of hiPSC, phPFC, hPFC, PYS and Epi derivatives at sampling days. (c) Bubble plot representing the frequency of expression and average expression of key lineage-associated genes in identified clusters in Fig.3g. (d) Heatmap showing the cosine similarity of Epi, pPGC, PSA-Epi and AMA-Epi in gastruloids, calculated from highly variable features. (e) Heatmap of DEGs of Epi, pPGC, PSA-Epi and AMA-Epi (middle panel). The representative marker genes (left panel) and related GO enrichment terms (right panel) were shown.

**Extended Data Fig. 11.**
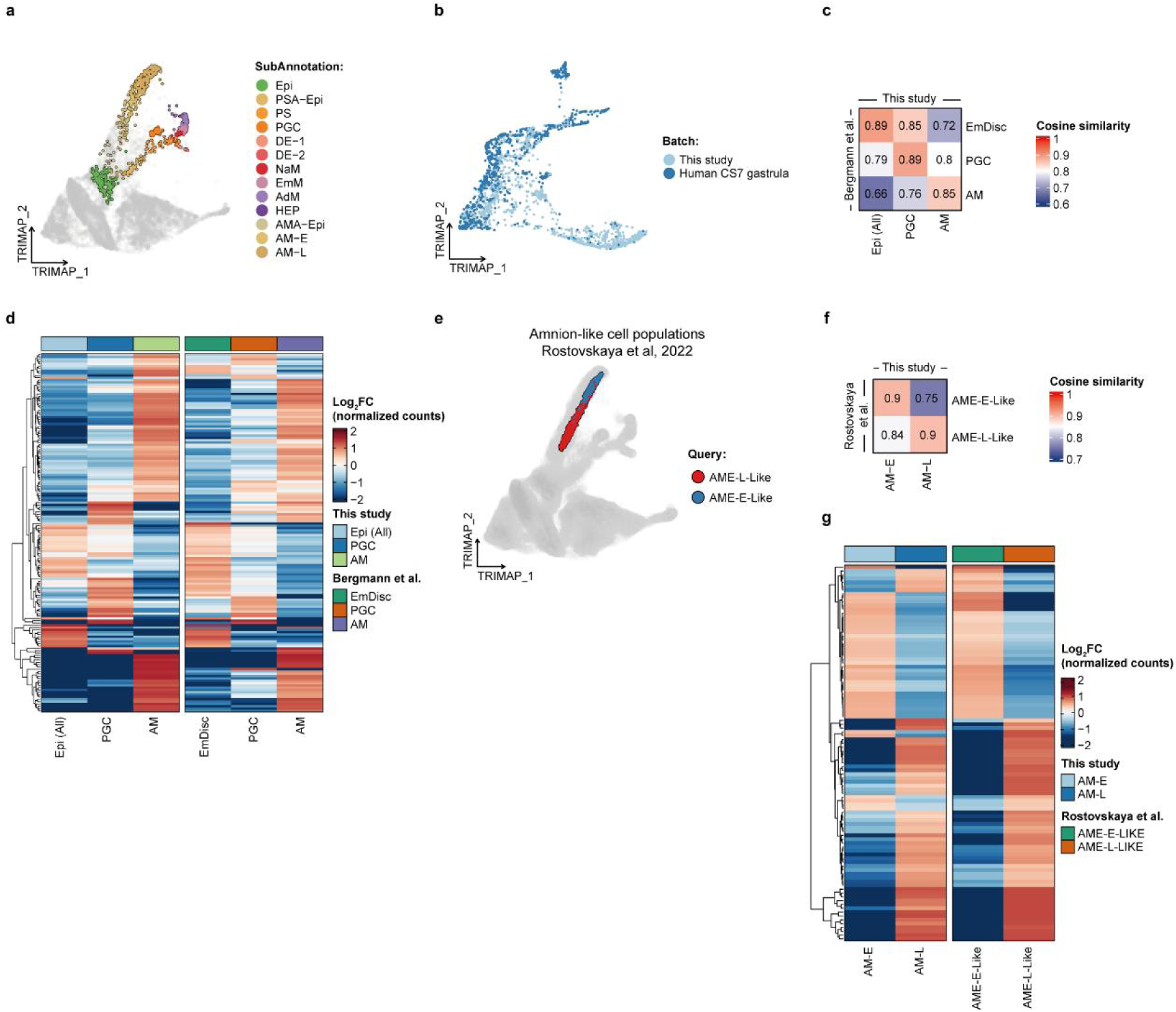
Similarity analysis of derivatives of epiblast in gastruloids. (a) TriMap showing the distribution of cells integrated with human CS7 gastrula dataset^13^ with highlighted borders in Fig. 3g. (b) TriMap showing the distribution of human CS7 gastrula dataset^13^ and the data of this study. (c) Heatmap showing the cosine similarity of AM, PGC and embryonic disc (EmDisc) in marmoset datasets versus AM, PGC and all kinds of Epi in gastruloids, calculated from shared DEGs. (d) Heatmap of the average expression of shared DEGs used in (c). (e) TriMap projection of the *in vitro* amnion-like cell populations dataset ^27^ onto amnion cells (namely AMA-Epi, AM-E and AM-L) of this study (grey). (f) Heatmap showing the cosine similarity of early and late amnion-like cells versus AM-E and AM-L in gastruloids, calculated from shared DEGs. (g) Heatmap of the average expression of shared DEGs used in (f).

**Extended Data Fig. 12.**
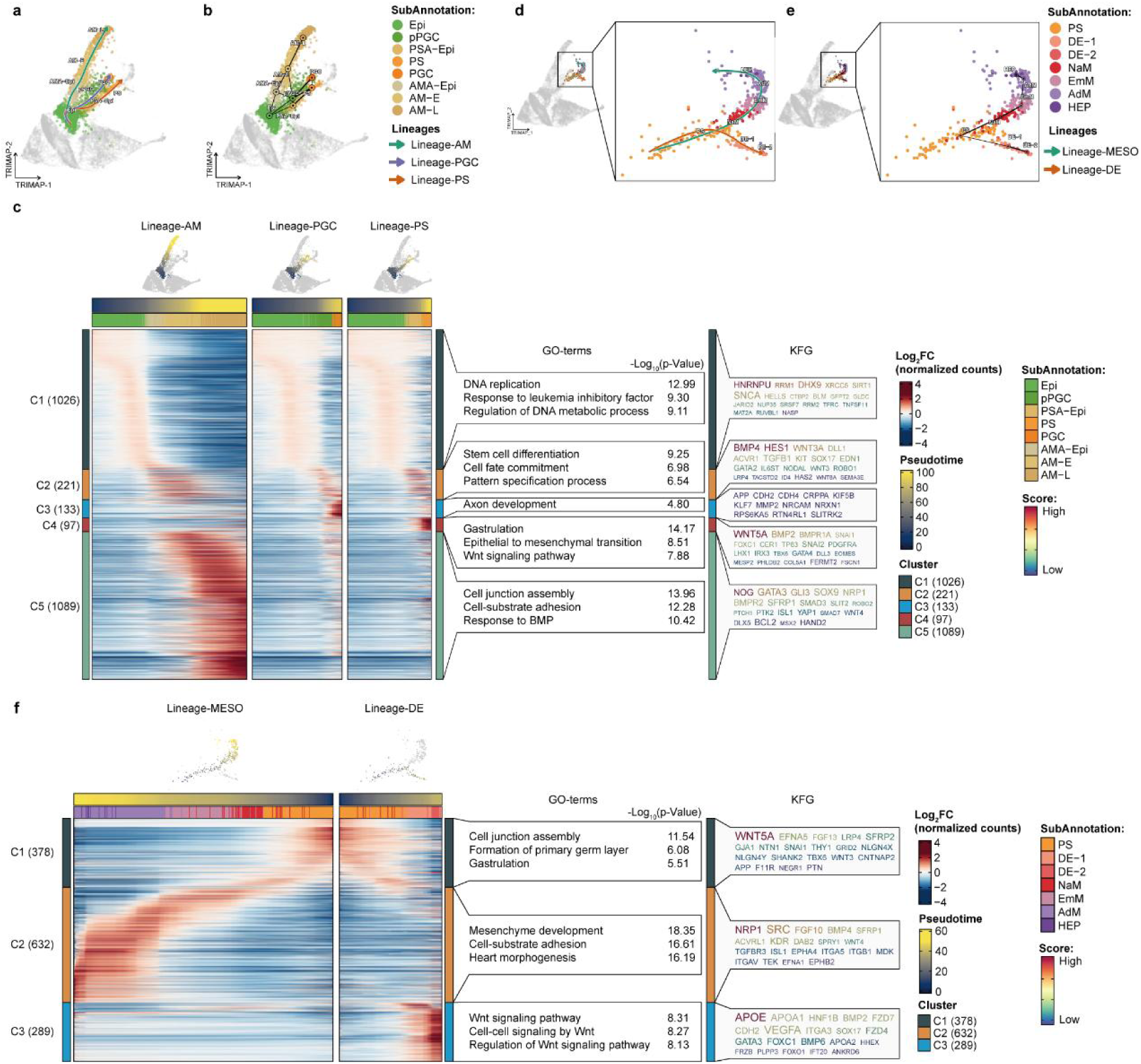
scRNA-seq analysis of the epiblast development trajectories. (a) TriMap showing cell trajectories inferred by Slingshot where cells underwent transitions from Epi to AM-L, from Epi to PGC and from Epi to PS. (b) PAGA analysis of Epi derivatives (excluding mesoderm and DE) overlaid on TriMap. The threshold for connection of cell types was set to 0.12. (c) Heatmap showing expression pattern dynamics (left panel) alongside the developmental trajectories from Epi to AM-L, from Epi to PGC and from Epi to PS with enriched GO enrichment terms (middle panel) and KFGs (right panel). TriMaps colored by pseudotime were annotated above the corresponding trajectories. (d) TriMap showing cell trajectories inferred by Slingshot where cells underwent transitions from PS to HEP and to DE-2. (e) PAGA analysis of PS derivatives overlaid on TriMap. The threshold for connection of cell types was set to 0.3. (f) Heatmap showing expression pattern dynamics (left panel) alongside the developmental trajectories from PS to HEP and to DE-2 with enriched pathways (middle panel) and KFGs (right panel). TriMaps colored by pseudotime were annotated above the corresponding trajectories.

**Extended Data Fig. 13.**
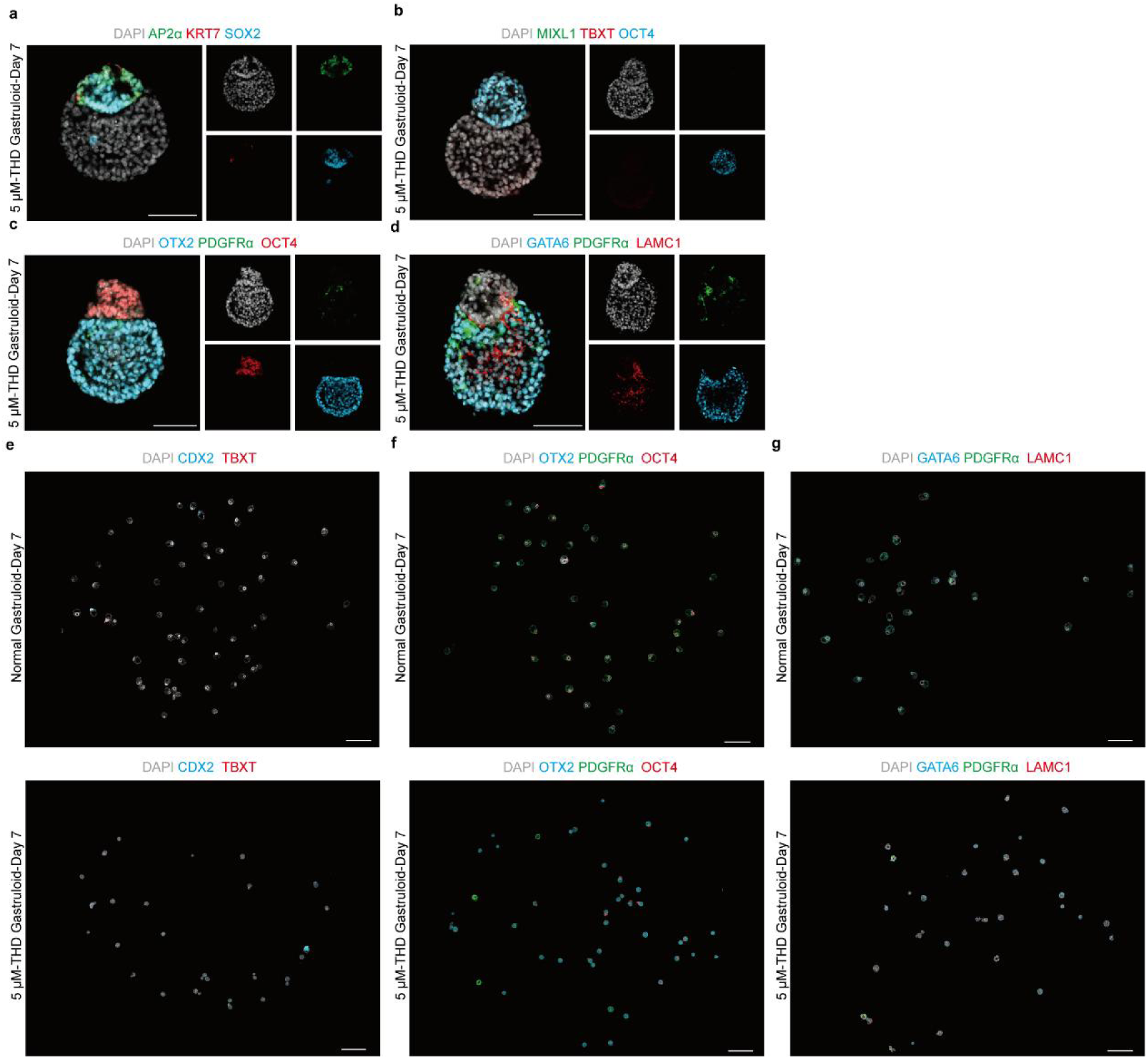
Morphology characterization of the gastruloids following THD treatment. (a) IF staining for epiblast marker SOX2 (blue), amnion cell marker AP2α (green) and KRT7 (red) of the aggregation (based on hiPSC) sustained with 5 μM THD on day 7. Nuclei were counterstained with DAPI (grey). Scale bar, 100 μm. (b) IF staining for epiblast marker OCT4 (blue), primitive streak marker TBXT (red) and mesoderm marker MIXL1 (green) of the aggregation (based on hiPSC) sustained with 5 μM THD on day 7. Nuclei were counterstained with DAPI (grey). Scale bar, 100 μm. (c) IF staining for epiblast marker OCT4 (red), hypoblast and anterior visceral endoderm marker OTX2 (blue) and secondary yolk sac marker PDGFRα (green) of the aggregation (based on hiPSC) sustained with 5 μM THD on day 7. Nuclei were counterstained with DAPI (grey). Scale bar, 100 μm. (d) IF staining for hypoblast marker GATA6 (blue), secondary yolk sac marker PDGFRα (green) and extraembryonic mesoderm marker LAMC1 (red) of the aggregation (based on hiPSC) sustained with 5 μM THD on day 7. Nuclei were counterstained with DAPI (grey). Scale bar, 100 μm. (e) Whole-mount IF staining for amnion marker CDX2 (blue) and primitive streak marker TBXT (red) of human gastruloids (upper) and the aggregates sustained with 5 μM THD (lower) (based on hiPSC) on day 7. Nuclei were counterstained with DAPI (grey). Scale bar, 1000 μm. (f) Whole-mount IF staining for epiblast marker OCT4 (red), hypoblast and anterior visceral endoderm marker OTX2 (blue) and secondary yolk sac marker PDGFRα (green) of human gastruloids (upper) and the aggregates sustained with 5 μM THD (lower) (based on hiPSC) on day 7. Nuclei were counterstained with DAPI (grey). Scale bar, 1000 μm. (g) Whole-mount IF staining for hypoblast marker GATA6 (blue), secondary yolk sac marker PDGFRα (green) and extraembryonic mesoderm marker LAMC1 (red) of human gastruloids (upper) and the aggregates sustained with 5 μM-THD (lower) (based on hiPSC) on day 7. Nuclei were counterstained with DAPI (grey). Scale bar, 1000 μm.

**Extended Data Fig. 14.**
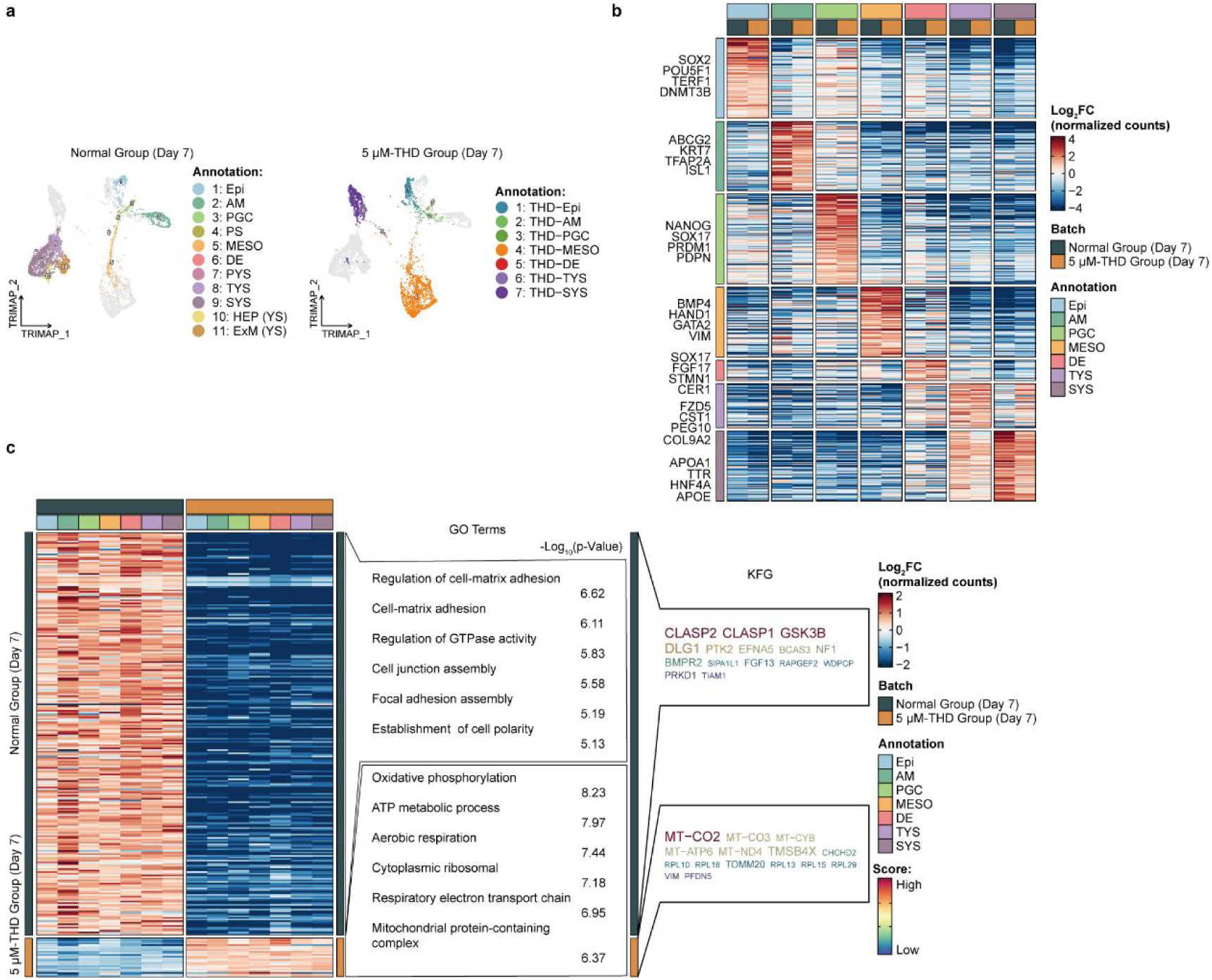
Transcriptional features of the gastruloids following THD treatment. (a) TriMaps showing cell types of gastruloids with 5 μM THD treatment integrated with the normal group on day 7 split by whether treated with THD. (b) Heatmap of the average expression pattern of DEGs conserved after THD treatment (right panel) with representative marker genes (left panel). (c) Heatmap of the average expression pattern of DEGs that produced changes in all cell types by THD treatment (left panel). The related GO enrichment terms (middle panel) and KFGs (right panel) were shown.

**Extended Data Fig. 15.**
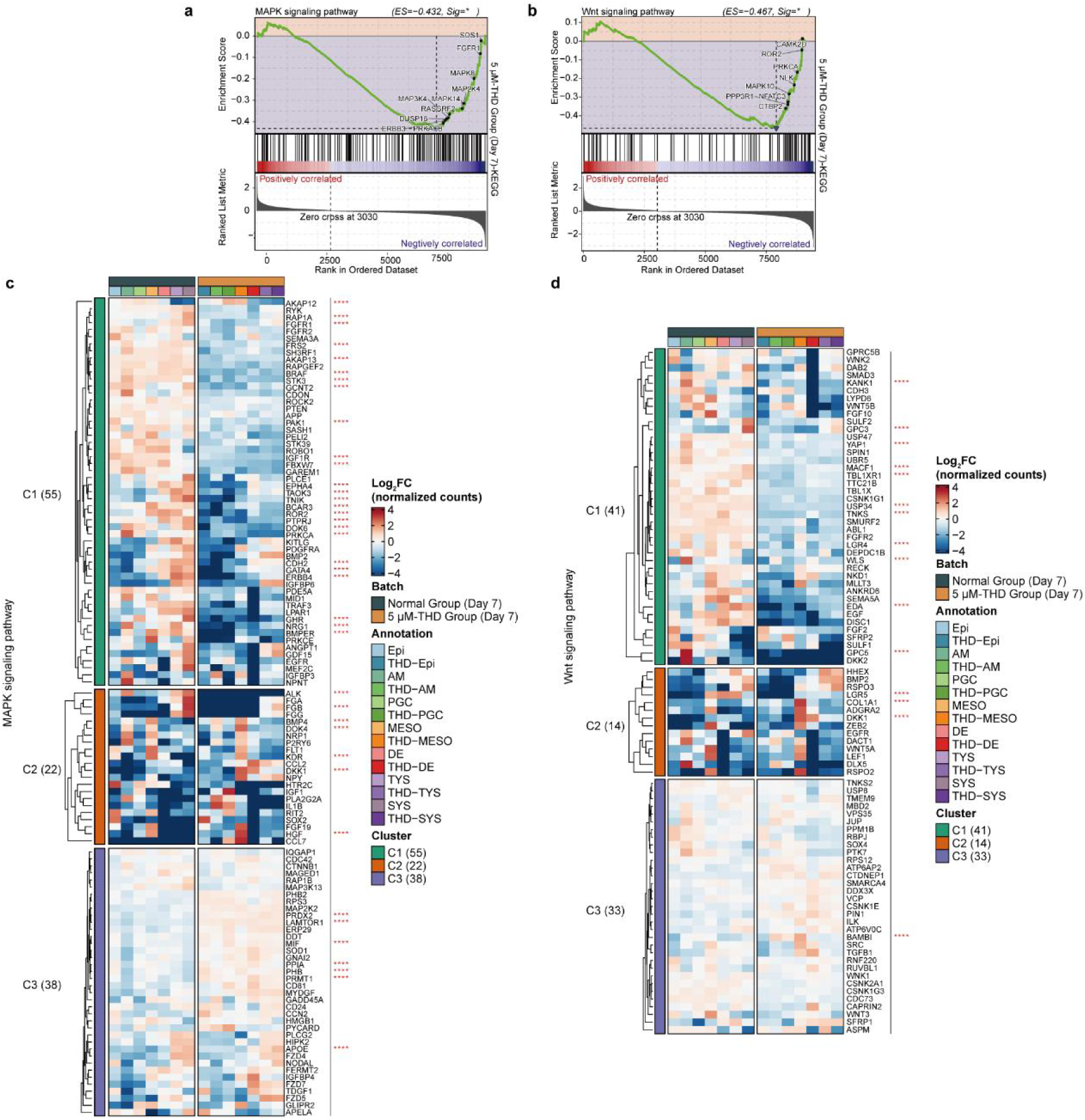
Analysis of MAPK and Wnt signaling pathways of the gastruloids following THD treatment. (a-b) GSEA plots showing differences in MAPK (a) and Wnt (b) signaling pathways after THD treatment compared with normal group. (c-d) Heatmaps showing the expression of genes about positive regulation of MAPK (ref: GO:0043410) (c) and Wnt (ref: GO:0030177) (d) signaling pathways with and without THD treatment between the corresponding cell types. C1 marked genes that were broadly decreased since THD treatment and C2 marked genes with larger fold changes between cell types than C3. P-values were marked in red.

## Materials and Methods

### Cell source and Ethical approval

The development of embryo-like model generated in this study lacks trophoblast, thus cannot form placenta. Therefore, our model does not have any potential to form human organismal morphology. Furthermore, all hESC and hiPSC studies were carried out after the approval from the ethical committee of Nanjing Medical University (No. 2019935) and adhered to the Ethical Guidelines for Human Embryonic Stem Cell Research issued by the Ministry of Science and Technology and the Ministry of Health of the People’s Republic of China in 2003 as well as the 2021 ISSCR Guidelines for Stem Cell Research and Clinical Translation.

### Culture medium Essential 8 medium

Essential 8 medium consisted of essential 8 basal medium (Gibco, A1517001) supplemented with 50x essential 8 supplement.

### Induction of 3D gastruloids first-stage medium

Induction of 3D gastruloids first-stage medium consisted of a 2:1:1 mixture of G-1^TM^ plus medium (Vitrolife, 10128), Essential 8 medium and RACL medium (PRMI 1640 medium (Gibco, 61870036), supplemented with 1% B27 supplement (Gibco, 12587-010), 0.1 mM 2-mercaptoethanol (Gibco, 21985023), 0.1 mM MEM non-essential amino acids (Gibco, 11140050), 3 μM CHIR99021 (MedChemExpress,HY-10071), 100 ng/mL Activin A (R&D system, 338-AC) and 20 ng/mL human LIF (R&D system, 225-SC).

### Induction of 3D gastruloids second-stage medium

Induction of 3D gastruloids second-stage medium consisted of a 2:1:1 mixture of G-2^TM^ plus medium (Vitrolife, 10132), Essential 8 medium and EBB medium (Essential 6^TM^ medium (Gibco, A1516401) supplemented with 0.1 mM GlutaMax (Gibco, 35050-061), 0.1 mM 2-mercaptoethanol, 0.1 mM MEM non-essential amino acids, 20 ng/mL BMP4 (R&D system, 314-BP) and 10 ng/mL bFGF (R&D system, 3718-FB).

### Culture of human pluripotent stem cells

The human PSCs culturing conditions were as follows: 37 °C, 5% CO_2_ and saturated humidity. DYR0100 hiPSC line and H1 hESC were maintained in a standard feeder-free culture using Essential 8 medium and plated in the 6-well plates coated with matrigal matrix (Corning, 354230) per the manufacturer’s instructions. DYR0100 hiPSCs were used before passage 45 and H1 hESCs were used before passage 55.

### Induction of 3D gastruloids

To induct gastruloids, the following conditions were used: a temperature of 37 °C, 5% CO_2_ and saturated humidity. hPSCs were treated with TrypLE Express (Gibco, 12605028), at 37 °C for 3 minutes, followed by gentle mechanical dissociation using a pipette. After centrifugation, the cell pellet was resuspended in first-stage medium, supplemented with Y-27632 (10 µM, MedChemExpress, HY-10583).

Next, 7.5-9.0×10^4^ cells were mixed with 0.5 mL culture medium and seeded into one well of 24-well aggreWell^400^ culture plate pretreated with anti-adherence rinsing solution (Stem Cell Technologies, 07010) following the manufacturer’s instruction. The cells were allowed to form aggregates within the microwell for a period ranging from day 0 to 1 depending on the cell lines and their propensity for aggregation. During the culture process, the first-stage medium was refreshed every 24 hours, with half of the medium changed each time.

After 3 days, the first-stage medium was replaced with the second-stage medium. Between day 4 and 7 of culture, the concentration of BMP4 used was adjusted based on the overall situation of different cell lines. In this study, the DYR0100 hiPSC was cultured for 5 days and BMP4 concentration was increased to 100 ng/mL and then decreased to 75 ng/mL after 24 hours. For the H1 hESC, BMP4 concentration was increased to 75 ng/mL and then decreased to 50 ng/mL after 24 hours. After day 6, the medium was refreshed with half of the medium, while keeping BMP4 concentration unchanged.

### Immunofluorescence staining

The samples were fixed with 4% paraformaldehyde in phosphate-buffered saline for 20 min at room temperature, washed three times with DPBS (Corning, R21-031-CV) (permeabilized with 1% Triton X-100 (Sangon, A110694) all the time). After blocking with 5% BSA (Sigma-Aldrich, V900933) in DPBS for 4 hours at room temperature, samples were then incubated with primary antibody diluted in blocking buffer 24 hours at 4 °C. After primary antibody incubation, samples were washed three times with DPBS and incubated with fluorescence-conjugated secondary antibodies diluted in blocking buffer at temperature for 4 hours. Nuclei were stained with DAPI (Gibco, D1306) at 1 μg/mL. Samples were washed three times with DPBS and waited for clarity.

The primary antibodies included: mouse anti-OCT4 antibody (Santa Cruz Biotechnology, Sc-5279; 1:100), rabbit anti-OCT4 antibody (Abcam, ab181557; 1:200), rabbit anti-SOX2 antibody (Abcam, ab92494; 1:200), goat anti-SOX2 antibody (R&D system, AF2018; 1:200), rabbit anti-NANOG antibody (Cell signaling tech, 4903S; 1:200), goat anti-NANOG antibody (R&D system, AF1997; 1:200), goat anti-GATA6 antibody (R&D system, AF1700; 1:500), goat anti-SOX17 antibody (R&D system, AF1924; 1:500), goat anti-OTX2 antibody (R&D system, AF1979; 1:200), rabbit anti-CDX2 antibody (Cell signaling tech, 12306; 1:200), rabbit anti-PDGFRa antibody (Abcam, ab203491; 1:200), rabbit anti-LHX1 antibody (Developmental Studies Hybridoma Bank, 4F2; 1:200), mouse anti-LAMC1 antibody (R&D system, MAB2139; 1:200), goat anti-TBXT antibody (R&D system, AF2085; 1:200), mouse anti-EZRIN antibody (Sigma-Aldrich, E8897; 1:200), rabbit anti-ISL1 antibody (Invitrogen, PA5-27789; 1:200), rabbit anti-EOMES antibody (Sigma-Aldrich, HPA028896; 1:200), rabbit anti-MXIL1 antibody (Proteintech, 22772-1-AP; 1:200), mouse anti-AP2γ antibody (Santa Cruz Biotechnology, SC-12762; 1:100), rabbit anti-BLIMP1 antibody (Cell signaling tech, 9115; 1:200), mouse anti-AP2α antibody (Santa Cruz Biotechnology, SC-12726; 1:100), rabbit anti-KRT7 antibody (Zsbio, ZA-0573; 1:200), goat anti-VIM antibody (R&D system, AF2105; 1:200).

The secondary antibodies (Invitrogen) included Alexa Fluor 488 donkey anti-mouse antibody (A-21202; 1:200), Alexa Fluor 488 donkey anti-rabbit antibody (A-21206; 1:200), Alexa Fluor 488 donkey anti-goat antibody (A-11055; 1:200), Alexa Fluor 555 donkey anti-mouse antibody (A-31570; 1:200), Alexa Fluor 555 donkey anti-rabbit antibody (A-31572; 1:200), Alexa Fluor 555 donkey anti-goat antibody (A-21432; 1:200), Alexa Fluor 647 donkey anti-mouse antibody (A-31571; 1:200), Alexa Fluor 647 donkey anti-rabbit antibody (A-32795; 1:200), Alexa Fluor 647 donkey anti-goat antibody (A-21447; 1:200).

### Sample clarity

#### Clearing solution preparation

20 g Histondenz^TM^ (Sigma-Aldrich, D2158) dissolved in 15 mL 0.2x PBS buffer, long-time storage at room temperature.

#### Clarity and fixation

A total of 45 μL of Clearing Solution was carefully added to the CoverWell™ Incubation Chamber Gaskets (Thermo Fisher Scientific, C18155). The cleaned samples, post-incubation with the secondary antibody, were then transferred to these incubation chamber gaskets. A thorough mixing of the samples within the Clearing Solution was achieved using a mouth pipette. The coverslip surface was gently covered from the side of the CoverWell™ Incubation Chamber Gaskets, taking care to avoid excessive foaming. Finally, the fixed sample was stored in a dark location until it was fully cleared.

#### Microscopy and image analysis

The brightfield image were taken with an inverted fluorescence microscope TE2000-s (Nikon) equipped with an AxioCam HRc camera (Zeiss) and another inverted Microscope Eclipse Ti2-U (Nikon). Confocal immunofluorescence images of human gastruloid were acquired with Zeiss LSM 800 confocal microscopes. Images were processed by ZEN (Zeiss).

For cell counting and volume measurement, Imaris (v8.0.0) software was used. Cell count parameters were set for size and fluorescence strength of voxels and then overall cell count data was obtained for each image using Imaris’s total number of spot function. Volume statistics by setting parameters was obtained for each image using Imaris’s volume function. Spots were drawn using the internal algorithm, using an estimated XY size of 8-10 μm, a quality threshold of 4 and background subtraction.

#### Thalidomide application

Set thalidomide (MedChemExpress, HY-14658) experimental groups (5 μM,10 μM) and normal, DMSO-control ((Sigma-Aldrich, D2650), corresponding volume to 10 μM thalidomide solvent) control group. Thalidomide was persistently added during gastruloid induction, with samples taken and validated seven days post-administration.

#### Induction proportion statistics

Experimental data from ten distinct batches of both H1 hESC and DRY0100 hiPSC were utilized. In each batch, over 20 gastruloids were randomly sampled and stained for further analysis. Subsequently, data were collected and analyzed in accordance with established standards.

### Chromium next GEM single cell RNA 3’library construction

We adhered strictly to the instructions provided in the 10x Genomics user guide for all steps of the process. The Chromium Next GEM Single Cell 3ʹ GEM, Library & Gel Bead Kit v3.1 (10x Genomics) was utilized for our experiments. Briefly, we prepared a mixture of all single-cell suspensions and reagents, and loaded this into the chip for GEM generation, facilitated by the Chromium Controller. We conducted GEM-RT incubation within the droplets and proceeded to recover single-strand cDNA through demulsification and dynabeads cleanup. Post-PCR, we applied SPRIselect to the pre-amplified cDNA product and carried it forward to library construction. We concluded by quantifying the libraries, undertaking quality control, and sequencing on an Illumina NovaSeq 6000 sequencer at the Nanjing Jiangbei New Area Biopharmaceutical public service platform in Nanjing, China.

### 10x Genomics data pre-processing

With the usage of the program 10x Genomics Cell Ranger (v5.0.0), sequencing data was processed^1^ to align, filter and count unique molecular identifiers (UMIs) per sample using the “--include-introns” parameter and then mapped to reference genome (refdata-gex-GRCh38-and-mm10-2020-A). Velocyto (v0.17.17)^2^ was used to process the output files and create loom files for the following RNA velocity analysis.

### Quality control and filtering

We utilized the scDblFinder^3^ (v1.10.0) algorithm per sample with default parameters to predict potential doublets. All cells that were flagged as doublets were excluded from the further downstream biological analysis. Cells with the number of detected genes (nFeature_RNA) above 3000 and detected transcripts (nCount_RNA) above 5,000 were retained to exclude cells with low complexity. We applied a high and low median absolute deviation (MAD) thresholds to exclude outlier cells. For log10_nCount and log10_nFeature, we determined outliers based on 2.5MADs looked for at both tails. For featcount_dist (distance to expected ratio of log10_nCount and log10_nFeature), we determined outliers based on 2.5MADs looked for only at the lower tail. To further generate high-quality data, cells were excluded from all samples if they had UMI counts for human mitochondrial genes exceeding 20%, UMI counts for ribosomal genes exceeding 50%, or a ratio of ribosomal and mitochondrial genes exceeding 50.

### Integrating time points data

We integrated the data from different induction time points and preprocessed it using Seurat ^4^ (v4.1.0) R package. After log-normalization, the default vst approach was used for each time point to find 3,000 highly variable genes. The *SelectIntegrationFeatures* function was used to choose the 3,000 most variable genes across all time points to create a set of characteristics appropriate for integration. The normalized expression levels were z-transformed, and the dimension was reduced based on Principal Component Analysis (PCA).

Scanorama is an integration approach that has been shown to be sensitive to subtle temporal changes within the same cell lineage. We used Scanorama with the default parameters for integrating cells at every time point^5^. For the content that data of this study participated, we used a two-dimensional representation of the data obtained by performing TriMap dimensionality reduction, which better preserves the global structure of the dataset. To perform the nonlinear reduction, the *TRIMAP* function of the trimap^6^ (v1.1.4) package was used on the Scanorama matrix. Cell cycle scores and phases were counted using the *CellCycleScoring* function and the cc.genes.updated-2019 cell cycle genes. TriMaps were performed with regression of S and G2M cell cycle scores using the vars.to.regress parameter in the *ScaleData* function. Integrated datasets were clustered based on the SNN graph using the *FindClusters* function.

### Differential gene expression analysis and enrichment analysis

To identify genes that are expressed differently in a specific cell type, we used the *FindMarkers* and *FindConservedMarkers* function of the Seurat pipeline to compare that cell type with those the others. The Wilcoxon test was used for comparison and the resulting p-value was adjusted using Benjamini-Hochberg (BH) method. Enrichment with differentially expressed genes (DEGs) filtered for each cell type (p_adjust<0.05, logfc.threshold>log2 (1.5)) was done using the *enricher* function of the clusterProfiler^7^ (v4.6.0) package with parameters minGSSize=10 and maxGSSize=500. The Biological Process (BP), Molecular Function (MF), and Cellular Component (CC) annotation in the org.Hs.eg.db (v3.15.0) package was used for gene ontology (GO) enrichment. We used the Kyoto Encyclopedia of Genes and Genomes (KEGG) database for gene set enrichment analysis (GSEA) to focus on changes in specified pathways. Genes that are located in highly enriched GO terms and have a high frequency of occurrence are considered as Key Functional Genes (KFGs). We defined and calculated the functional gene score: if a gene *g* appears in a total of *n* enriched terms, where the adjusted p-value of the *i* the term is *p_i_*, then the functional score of the gene *g* is:

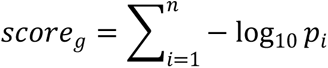

### PAGA and scVelo analysis

The scanpy (v1.9.1) implementation^8^ was used for Partition-based Graph Abstraction (PAGA) analysis^9^. The neighbor graph was constructed on the integrated Scanorama space using the *scanpy.pp.neighbors* function. Cell identity labels were used as partition categorical for the PAGA architecture. We used the *scanpy.tl.paga* with default settings to calculate the PAGA graph.

RNA velocities were analyzed using the scVelo^10^ (v0.2.4) package. We executed the *scvelo.pp.moments*, *scvelo.tl.velocity*, and *scvelo.tl.velocity_graph* functions to estimate velocity in default stochastic mode. The RNA velocity vector was projected onto TriMap using the *scvelo.pl.velocity_embedding_stream* function. In the part of hypoblast derivatives, we calculated genes with the step-changes in expression dynamics (n=18) according to the method of Barile et al.^11^ and removed them in the subsequent RNA velocity analysis.

### Slingshot and Dynamically Differential Gene (DDG) analysis

The published package slingshot (v2.6.0) was used to infer trajectories and align cells along a differentiation pseudotime^12^. The trajectories were determined using the slingshot wrapper function, TriMap dimensionality reduction, and cluster labeling as in Seurat objects. The slingshot trajectories are computed based on the cell types in the dataset with known start points or end points. After slingshot assigned pseudotime, 10,000 highly variable genes of each lineage were chosen as the candidates for DDGs. DDGs were filtered using the threshold: p_adjust<0.05, r.sq>0.1, and dev.expl>0.1. The fidelity of pseudotime was determined on the basis of gene clustering analysis. Furthermore, cells were arranged according to pseudotime and genes were ordered according to peak time of the fitted expression value. We divided clusters based on gene expression patterns and perform GO enrichment analysis on each cluster according to the method described above.

### Comparison of single-cell transcriptomic dataset among multiple datasets

We used the intersection of highly variable genes for projection between datasets based on k-nearest neighbors (KNN) algorithm, with the target number of neighbors set to 30. To confirm the identity of specific cell types in gastruloids, the human CS7 gastrula scRNA-seq dataset was integrated with the data generated in this study using fastMNN^13^ integration standard workflow (Fig. 2h, i, 3j, k, Extended Data Fig. 6h, i, 11a, b). In the epiblast derivative part, due to the diversity of cell types and the close numbers of cells between datasets, we used 3000 highly variable genes from the human CS7 gastrula scRNA-seq dataset for integration. In the hypoblast derivatives section, to highlight the features of visceral endoderm (VE) and yolk sac endoderm (YSE) in CS7 gastrula, we used DEGs in the human CS7 gastrula scRNA-seq dataset for integration. For cross-species comparison, we converted marmoset genes to human homologous gene with the biomaRt package. Cosine similarity with the marmoset gastrulation datasets were calculated on the basis of the intersection genes across DEGs of datasets.

The dataset of pluripotent founder cell (hPFC) published as reference is available at GEO under accession code: GSE126022. The dataset of human CS7 gastrula used as reference are available from ArrayExpress under accession code E-MTAB-9388 and the processed data can be downloaded from http://www.human-gastrula.net. The dataset of *in vitro* cultured (IVC) human embryos used as reference is available at the Gene Expression Omnibus (GEO) under accession number GSE136447. The dataset of IVC human post-implantation embryos is available at the ArrayExpress database under accession code: E-MTAB-8060. The dataset of human pre-implantation embryos as reference is available at the ArrayExpress database under accession code: E-MTAB-3929 and cell type annotations proposed by Stirparo et al.^14^ were used. The dataset of marmoset gastrulation in utero used as reference is available at the ArrayExpress under accession numbers E-MTAB-9367. The dataset of in vitro amnion-like cell populations as reference is available at GEO under accession code: GSE179309.

