## Supplemental tables and videos for "Establishment of a novel non-integrated human pluripotent stem cell-based gastruloid model": SI guide-final.docx

**Supplementary Table 1 | Differential expressed genes list.**

A list of all differential expressed genes (DEGs) and the corresponding Gene ontology (GO) enrichment results in this study, including: DEGs of 12 major cell types; DEGs of epiblast (Epi), precursor of primordial germ cell (pPGC), primitive streak anlage-epiblast (PSA-Epi) and amnion anlage-epiblast (AMA-Epi); conserved and disturbed DEGs among cell types with and without thalidomide (THD) treatment.

**Supplementary Table 2 | GO enrichment results of differential developmental genes.**

A list of GO enrichment results of all differential developmental genes (DDGs) in this study, including: DDGs along trajectories from human pluripotent founder cell (hPFC) to haemato-endothelial progenitor (HEP) and to extraembryonic mesoderm of secondary yolk sac (SYS-ExM); DDGs along trajectories from Epi to amnion-Late (AM-L), from Epi to PGC and from Epi to primitive streak (PS); DDGs along trajectories from PS to HEP and to definitive endoderm (DE).

**Supplementary Table 3 | Gene list used for cosine similarity calculation.**

A list of shared DEGs of datasets and this study used for cosine similarity analysis, including: visceral endoderm (VE), secondary yolk sac (SYS) and ExM/Stalk in marmoset datasets versus VE or anterior visceral endoderm (AVE), the parietal endoderm of SYS (SYS-PE) and ExM in gastruloids; AM, PGC and embryonic disc (EmDisc) in marmoset datasets versus AM, PGC and all kinds of Epi in gastruloids; early and late amnion-like cells versus AM-E and AM-L in gastruloids.

**Supplementary Movie 1 | A 3D video captured by a light sheet microscope of the human gastruloid on day 7.**

A 3D video of the human gastruloid on day 7 shows 3D shape of the day 7 gastruloid including the amniotic cavity, embryonic disc, and secondary yolk sac. Immunofluorescence staining for epiblast marker OCT4 (red), amnion marker ISL1 (green) and hypoblast marker GATA6 (blue).

**Supplementary Movie 2 | A 3D video captured by a light sheet microscope of the human gastruloid on day 7 demonstrating the anterior visceral endoderm.**

A 3D video of the human gastruloid on day 7 shows 3D shape of the day 7 gastruloid including anterior visceral endoderm, embryonic disc, and secondary yolk sac. Immunofluorescence staining for epiblast marker OCT4 (red), anterior visceral endoderm marker OTX2 (blue) and secondary yolk sac marker PDGFRα (green).

**Supplementary Movie 3 | A 3D video captured by a light sheet microscope of the human gastruloid on day 7 demonstrating the gastrulation.**

A 3D video of the human gastruloid on day 7 shows 3D shape of the day 7 gastruloid including primitive streak, embryonic disc, and secondary yolk sac. Immunofluorescence staining for epiblast marker OCT4 (blue), primitive streak marker TBXT (red) and endoderm marker EOMES (green). Nuclei were counterstained with DAPI (gray).
